# Cell atlas of the regenerating human liver after portal vein embolization

**DOI:** 10.1101/2021.06.03.444016

**Authors:** Agnieska Brazovskaja, Tomás Gomes, Christiane Körner, Zhisong He, Theresa Schaffer, Julian Connor Eckel, René Hänsel, Malgorzata Santel, Timm Denecke, Michael Dannemann, Mario Brosch, Jochen Hampe, Daniel Seehofer, Georg Damm, J. Gray Camp, Barbara Treutlein

**Affiliations:** Max Planck Institute for Evolutionary Anthropology, Leipzig, Germany; Department of Biosystems Science and Engineering, ETH Zürich, Basel, Switzerland; Department of Hepatobiliary Surgery and Visceral Transplantation, Leipzig University, Leipzig, Germany; Rudolf-Schönheimer Institute for Biochemistry, Faculty of Medicine, Leipzig University, Leipzig, Germany; Institute for Medical Informatics, Statistics and Epidemiology (IMISE), Leipzig University, Leipzig, Germany; Department of Diagnostic and Interventional Radiology, Leipzig University, Leipzig, Germany; Institute of Genomics, University of Tartu, Tartu, Estonia; Medical Department 1, University Hospital Dresden, Technical University Dresden, Dresden, Germany; Center for Regenerative Therapies Dresden (CRTD), Technische Universität Dresden (TU Dresden), Dresden, Germany; Institute of Molecular and Clinical Ophthalmology Basel, Switzerland; Department of Ophthalmology, University of Basel, Switzerland

**Keywords:** liver, scRNA-seq, regeneration, human, portal vein embolization

## Abstract

The liver has the remarkable capacity to regenerate. In the clinic, this capacity can be induced by portal vein embolization (PVE), which redirects portal blood flow resulting in liver hypertrophy in locations with increased blood supply, and atrophy of embolized segments. Here we apply single-cell and single-nucleus transcriptomics on healthy, hypertrophied, and atrophied patient-derived liver samples to explore cell states in the liver during regeneration. We first establish an atlas of cell subtypes from the healthy human liver using fresh and frozen tissues, and then compare post-PVE samples with their reference counterparts. We find that PVE alters portal-central zonation of hepatocytes and endothelial cells. Embolization upregulates expression programs associated with development, cellular adhesion and inflammation across cell types. Analysis of interlineage crosstalk revealed key roles for immune cells in modulating regenerating tissue responses. Altogether, our data provides a rich resource for understanding homeostatic mechanisms arising during human liver regeneration and degeneration.

## Introduction

The human liver has extraordinary complexity on an architectural, cellular and functional level. This complexity allows the liver to execute diverse metabolic, immunological and detoxification functions across groups of coordinating cell types^1^. The liver is functionally organized into hexagonal lobules with the diverse cell type heterogeneity associated with patterns of blood flow^2^. Lobules are structured by blood vessels forming the portal triad, and converging at the center into the central vein. This portal-to-central axis was shown to have correlation with phenotypic and metabolic variation within hepatocytes referred to as zonation^3–5^, and liver sinusoidal endothelial cells (LSEC) show additional changes in adhesion molecules, and fenestrae size along this zonation axis in mouse and human^6–8^. Recent single-cell RNA-sequencing (scRNA-seq) studies have shed light into various aspects of liver biology in homeostasis^9–12^ and disease^13–15^. These datasets have provided unprecedented resolution of liver cell type heterogeneity, tissue structure and intercellular contacts, and have been powerfully used to reconstruct hepatocyte and LSEC zonation patterns. Despite these efforts, obtaining a representative sampling of all cell types in this tissue during physiological and pathophysiological conditions has been the object of intense research^16^.

In addition to its functional and structural complexity, the liver is able to restore itself and expand when injured or partially removed^17^. Liver regeneration comprises hypertrophy (cells increase in size) and hyperplasia (cells increase in number) as a compensation for lost tissue mass^18^. Mechanisms underlying cellular and molecular liver regeneration processes have been extensively studied using non-human models^19^ by applying commonly used experimental approaches such as partial hepatectomy or pharmacologically-induced liver damage^18^. While single-cell transcriptome sequencing has started to reveal cellular processes and states in mouse models of hepatectomy^20^, it is unclear how much of these insights can be transferred to humans. While hepatic resection has been used in the clinic to treat different liver diseases and has opened up opportunities to work on transplantation strategies^21^, temporally controlled experiments such as those performed in animal models following hepatectomy are not achievable in humans. This creates a gap in knowledge between animal and clinical studies on liver regeneration, and it is unknown how cell states undergo regeneration in the human liver.

A medical technique called portal vein embolization (PVE) provides an opportunity to gain insight into human liver regeneration. PVE is applied before surgery to avoid liver insufficiency in patients undergoing liver resection. This technique redirects blood flow to specific liver segments, resulting in a hypertrophic regenerated portion which functionally compensates the atrophied embolized portion to be surgically removed^21,22^. Here we apply single-cell and single-nucleus transcriptomics on healthy, hypertrophied, and atrophied patient-derived liver biopsies to explore cell states during this paradigm of human liver regeneration. The data are used to unravel cell types diversity and heterogeneity within cell types, biological pathways and intercellular communication responses during multiple physiological processes in the human liver.

## Results

### Fresh single-cell and frozen single-nucleus RNA-seq reveal major cell populations in adult healthy liver tissue samples

We performed single-cell and single-nucleus RNA-seq to generate a map of transcriptional profiles of the healthy human liver. We acquired human liver tissue samples assessed by pathologists as histologically healthy from patients undergoing liver resection (see Methods, Extended Data Table 1), and developed downstream processing protocols in two different ways (Fig. 1a). First, on the day of a surgery we obtained biopsies for an immediate experiment (referred to as fresh), and established a workflow to generate single-cell suspensions for parenchymal (hepatocytes) and non-parenchymal (cholangiocytes, endothelial, immune and mesenchymal cells) fractions (Extended Data Fig. 1a). Second, we used liquid nitrogen to snap freeze a portion of the biopsy, which we were able to store long term and use for preparation of single-nucleus suspensions (referred to as frozen) (Extended Data Fig. 2a,b). For frozen samples, single-nuclei were liberated from the tissue using Dounce homogenization and isolated using fluorescence activated cell sorting (FACS) with no additional fractionation as was performed in the fresh samples.

**Fig. 1.**
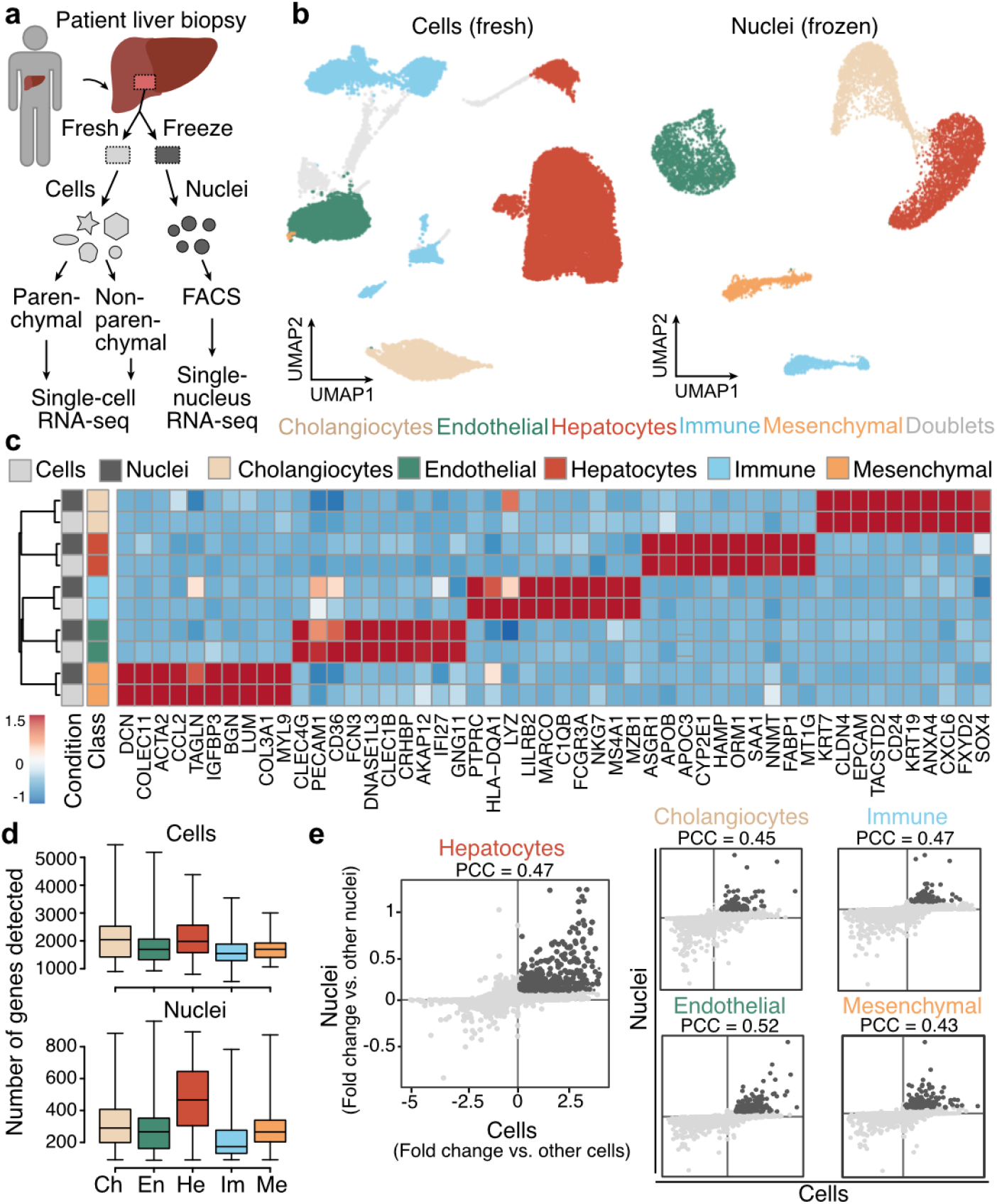
Fresh single-cell and frozen singlenucleus RNA-seq reveal major cell populations in adult healthy liver specimens. a, Single-cell and single-nucleus RNA-seq experiments were performed on patient-derived fresh (n=3) and frozen (n=3) liver tissues. Prior to single-cell RNA-seq, cells isolated from the fresh tissue were partitioned into parenchymal (hepatocytes) and nonparenchymal (other hepatic cell types) fractions (left). Prior to single-nucleus RNA-seq, nuclei from snap-frozen tissues were isolated using fluorescence activated cell sorting (right). b, UMAP plot of transcriptomes from fresh (left) and frozen (right) tissue datasets colored by major cell type (see Extended Data Fig. 1-2 for each donor representation). Gray represents potential doublets. c, Scaled expression of marker genes per major cell type across fresh and frozen tissue data (cells and nuclei are represented in rows, genes in columns). d, Number of detected genes per cell (top) and nucleus (bottom) across major cell types. Ch, cholangiocytes; En, endothelial cells; He, hepatocytes; Im, immune cells; Me, mesenchymal cells. e, Comparison of marker gene detection is shown for each major cell type in fresh and frozen samples (dark gray represents genes shared between both experimental setups; PCC, Pearson correlation coefficient).

We generated and analyzed transcriptomes of 21,000 cells and 9,400 nuclei from three fresh and three frozen samples with two samples in each condition derived from the same donor, respectively (see Methods, Extended Data Table 1). We analyzed fresh or frozen tissue datasets separately after integrating the cells from different donors^23,24^ (see Methods), and visualized cell heterogeneity using a uniform manifold approximation and projection (UMAP) embedding (Fig. 1b). Clustering revealed 5 previously reported major cell populations^9,12^ in both fresh and frozen tissue datasets representing hepatocyte, cholangiocyte, endothelium, mesenchyme and immune cells (Fig. 1b). Despite the lower number of genes detected per cell population in the frozen samples, cell type marker genes were consistently detected in both conditions, and differentially expressed genes between cell types were strongly correlated between fresh and frozen tissue datasets (Fig. 1c-e, Extended Data Fig. 1-2). The higher depth and sensitivity of the data from fresh whole-cell samples enabled a better resolution of cell subpopulations (Extended Data Fig. 1e-f). However, the unfractionated sampling using snRNA-seq provided a more representative survey of certain cell populations. For example, the proportion of mesenchymal cells in the frozen tissue dataset was over thirty times higher compared to fresh tissue data (Extended Data Tables 2 and 3). In addition, the enrichment of non-parenchymal immune populations from the fresh samples processing pipeline, revealed a larger fraction of immune cells and a more diverse set of subpopulations (Extended Data Fig. 1). We were able to reconstruct previously described zonation patterns in hepatocytes (Extended Data Fig. 3) and in LSEC (Extended Data Fig. 4) in both fresh and frozen tissue datasets with expression signatures shared between the experimental setups. Altogether, these data established strategies for generating single-cell and single-nucleus transcriptome atlases from fresh and frozen patient-derived liver samples.

### Transcriptional landscape of human liver cells after portal vein embolization

To investigate cellular processes specific to the regenerating human liver we obtained freshly resected liver tissue samples from six patients who underwent a preoperative medical procedure called portal embolization (PVE) (see Methods). During PVE the portal vein branching to the diseased part of the liver is blocked, or embolized with metal coils, and the future liver remnant receiving increased portal blood flow starts expand over time (Fig. 2a-b). On the resection day, we received two tissue samples from the same donor that we refer to as regenerating and embolized samples. We isolated parenchymal and non-parenchymal cell fractions from these samples in the same manner described above for the healthy condition, and performed scRNA-seq on each fraction. We integrated expression data from all three conditions (see Methods) and projected the data using UMAP (Fig. 2c, see Methods). In the combined dataset we identified hepatocytes, endothelial cells (EC), cholangiocytes, immune cells and mesenchymal cells as the major cell types (Fig. 2d and Extended Data Fig. 5), which were recovered in similar proportions from all donors and conditions (Fig. 2e and Extended Data Fig. 5c).

**Fig. 2.**
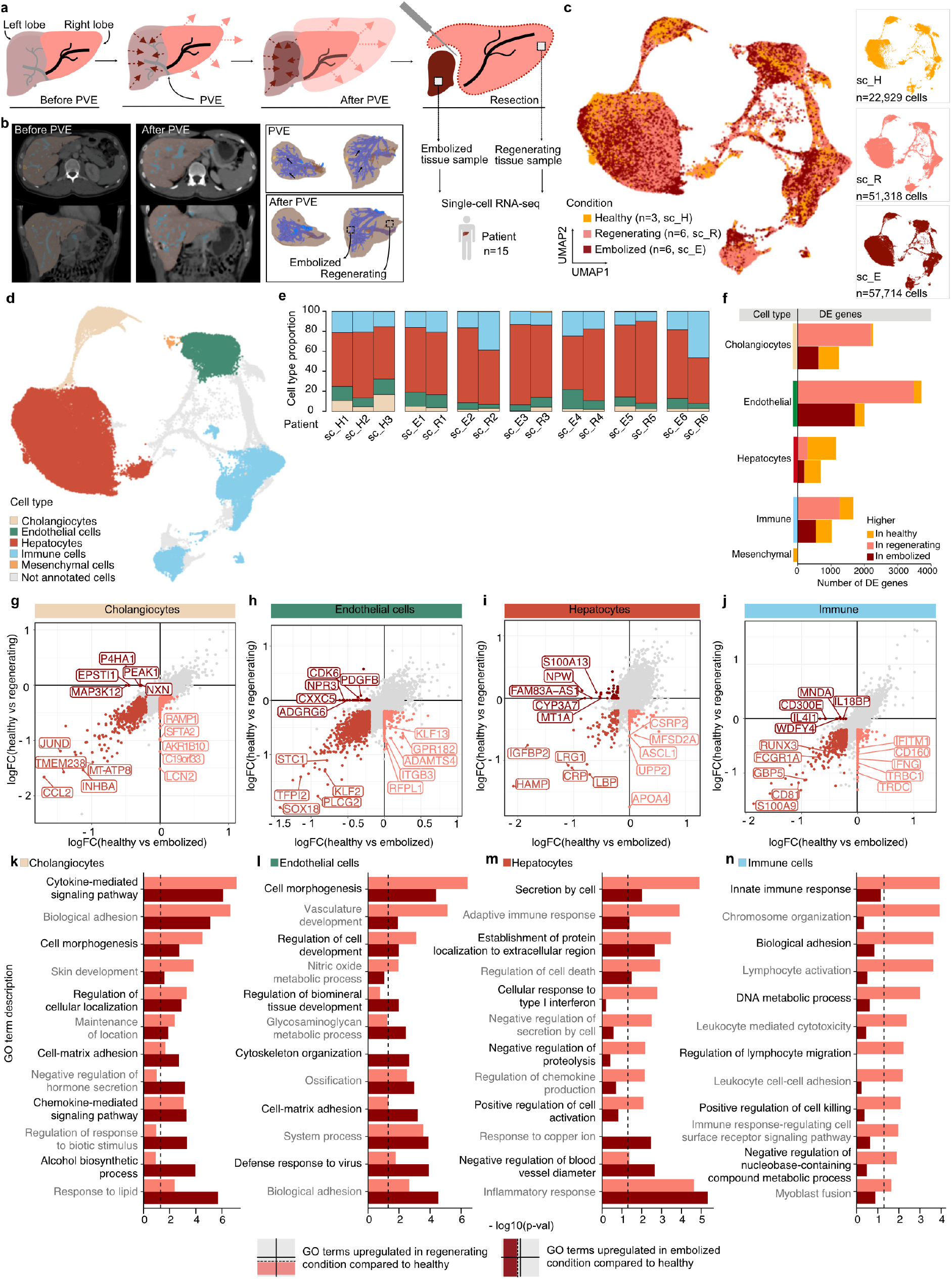
Transcriptional landscape of human liver cells after portal vein embolization. a, Schematic shows the portal vein embolization (PVE) procedure and tissue sampling for the scRNA-seq experiments. Portal vein branching toward diseased liver tissue (left lobe) prior to resection is blocked or embolized. Adjacent tissue with redirected blood flow (right lobe) expands over time. ScRNA-seq is performed on two samples derived from regenerating and embolized liver tissues on the day of liver resection. b, Computed tomography (CT) scans of pre-PVE (left) and post-PVE (middle) livers are shown along with 3D tissue reconstructions (right), respectively. c,d, UMAP plot of merged healthy (n=3), regenerating (n=6) and embolized (n=6) fresh liver samples, colored by condition (c) and major cell type (d). e, Major cell type proportion per donor and condition. f, Number of differentially expressed genes between healthy and regenerating or embolized samples per major cell type. g-j, Gene expression log fold change for each PVE condition compared to healthy (x-axis: embolized; y-axis: regenerating) for Cholangiocytes (g), Endothelial cells (h), Hepatocytes (i), and Immune cells (j). k-n, Enriched gene ontology terms for DE genes per major cell type, comparing regenerating or embolized to healthy. Top 6 terms were selected for each condition. Dashed line shows p-value=0.05.

For each cell type we compared gene expression levels in post-PVE conditions to their counterparts in healthy liver tissue. Differentially expressed (DE) genes in all cell types except hepatocytes were more often upregulated in both regenerating and embolized conditions (Fig. 2f, Extended Data Table 4). Such regeneration kinetics are consistent with previous results showing that hepatocytes are the first cell types entering the regeneration program after hepatectomy, and are followed by cell types responsible for tissue reorganization, including liver vasculature and bile ducts reestablishment^25,26^. Analysis of DE genes within cholangiocytes (Fig. 2g), ECs (Fig. 2h), hepatocytes (Fig. 2i) and immune cells (Fig. 2j), showed genes that were commonly upregulated in the cells from the regenerating and embolized tissue, as well as features that were specific to each condition compared to the healthy reference. For each major cell type there was a significant correlation pattern between both medical conditions compared to healthy, however the correlation was weakest for hepatocytes (Spearman’s rho = 0.48 for hepatocytes as compared to Spearman’s rho = 0.73 - 0.80 for ECs, cholangiocytes, immune cells) showing that responses between regenerating and embolized tissue are more divergent in this cell type.

DE genes revealed an enrichment for multiple expression programs in post-PVE samples (Fig. 2k-n). Gene ontology (GO) Terms related to development (“Cell morphogenesis” in cholangiocytes and “Vasculature development” in ECs) were present both in regenerating and embolized sections, with a stronger enrichment result in the former. GO terms associated with cellular adhesion pervade all major cell types, attesting to the considerable morphogenic changes induced by PVE. ECs and hepatocytes also revealed an upregulation of innate immune and inflammation programs (Fig. 2l,m), pathways with a critical role in initiating the regenerative process^27^. We also noted a specific enrichment for various pathways in immune cells in the regenerating tissue (Fig. 2n), an indication of their pivotal role in the process.

Among the transcripts that were shared between regenerating and embolized tissue hepatocytes were also genes shown to have a zone-specific expression (HAMP, CRP, IGFBP2, SAA1, SAA2; Fig. 2i). Additionally, we identified transcription factors (ASCL1, ATF5, CEBPA, FOXA3, IRF7, SREBF1) (Extended Data Table 4) that were DE between regenerating and healthy tissue, including Achaete–scute complex homolog-like 1 (ASCL1) which showed gestation-dependent higher expression in expanding livers of pregnant mice^28^, and CCAAT/enhancer-binding protein α (CEBPα) which exhibited antiproliferative functions and a potent regulator of the cell cycle exit^29^.

### Periportal-like hepatocytes predominate in post-PVE liver

Next we explored whether post-PVE hepatocytes show zonation of expression similar to hepatocytes in healthy tissues (Fig. 3a and 3b, Extended Data Fig. 3). UMAP embeddings of post-PVE hepatocytes revealed a portal-to-central signature gradient similar to healthy hepatocytes (Fig. 3c) with similar pseudo-zonated expression profiles of marker genes in all conditions (Fig. 3d). However, we observed a depletion of cells with a central-zone expression signature in post-PVE tissue hepatocytes and a corresponding enrichment in cells with a periportal-like signature (Fig. 3e). In order to evaluate these inferences on a spatial tissue level, we estimated the density of hepatocytes along the portal-central axes within patient tissue sections (see Methods). We confirmed a homogenous hepatocytes distribution in the healthy liver lobule. In contrast, lobules of PVE samples showed an increased hepatocyte density around the portal vein. While the lobule from the embolized tissue showed a pronounced decrease in pericentral hepatocytes density, the lobule from the regenerating tissue was more heterogeneous with an additional increase in hepatocytes density towards the periphery (Fig. 3f-h). This shift in cell density was accompanied by a change in the spatial arrangement of zonation markers (Extended Data Fig. 6). The central marker CYP3A4 was less often detected between the intermediate lobule area and the central vein, whereas the portal marker HAL was less often detected between the intermediate lobule border section and the acinar zones.

**Fig. 3.**
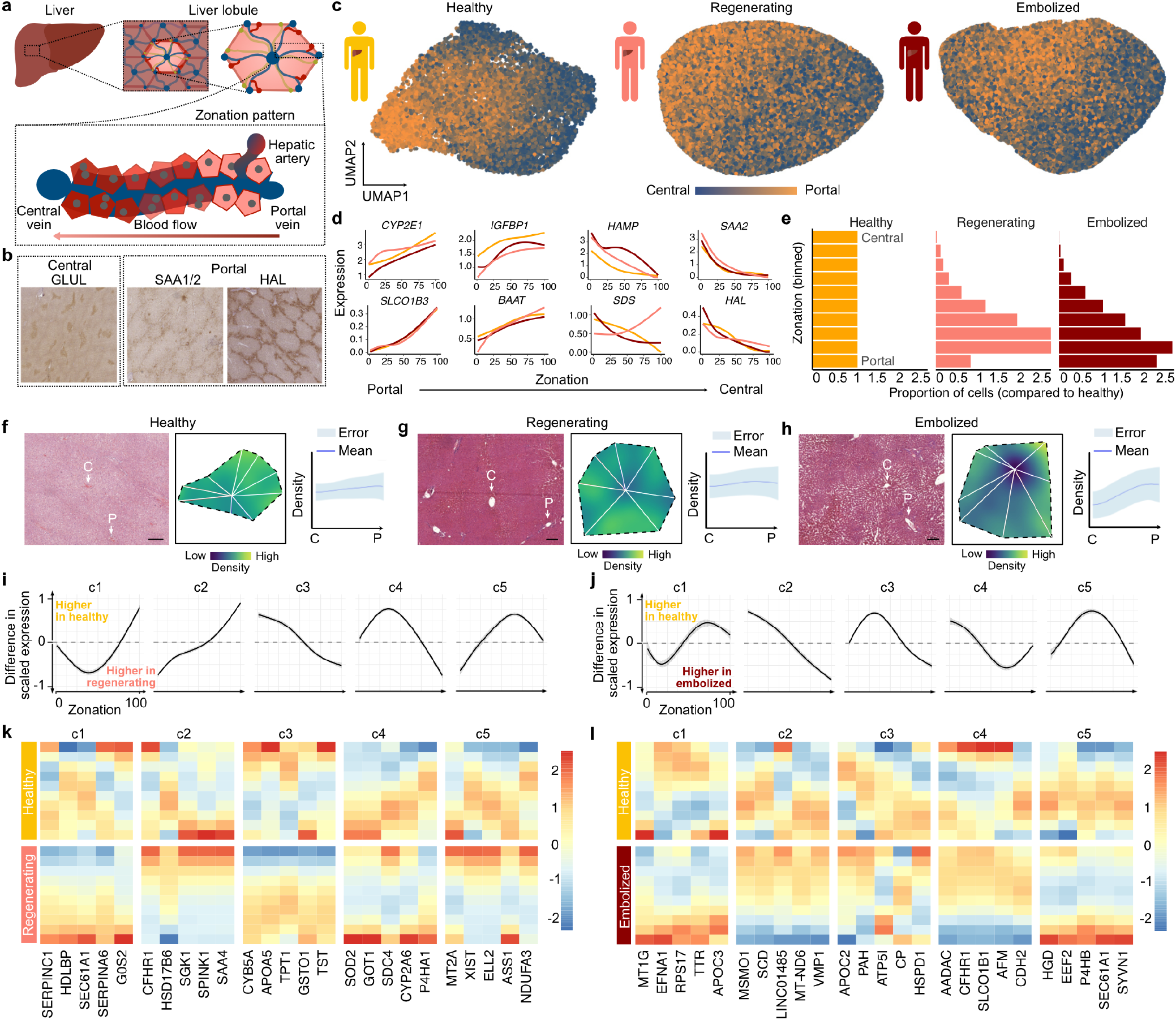
Spatial zonation patterns are altered in regenerating and embolized tissue hepatocytes. a, Schematic illustrating the structure of a liver lobule. b, DAB stainings of central and portal zone-specific protein expression within healthy tissue hepatocytes. c, Gene expression signature of zonation within healthy (right), regenerating (middle) and embolized (left) tissue hepatocytes. Cells in UMAP plots are colored based on the cumulative expression of central (blue) or portal (orange) zonation marker genes. d, Pseudozonation expression patterns of representative zonation marker genes for each medical condition. e, Histograms showing the proportion of hepatocytes in zonation bins across conditions. f-h, Representative H/E stainings are shown for healthy (f), regenerating (g) and embolized (h) tissue sections including a hepatocyte nuclei density analysis using a Kernel Density Estimation (KDE) algorithm presented as a corresponding heatmap of a representative liver lobule and quantification of the cell density along the portal-central axis (data show mean ± standard deviation, n = see Methods). i, Clusters of differential zonation patterns were identified between healthy and regenerating (i) or embolized (j) hepatocytes. k-l, Heatmaps show example genes with different zonation patterns between the healthy and regenerating (k) or embolized (l) tissue hepatocytes.

We next identified genes with diverging zonation patterns between healthy and regenerating or embolized tissue hepatocytes by correlating fitted expression values along zonation trajectory between healthy and post-PVE hepatocytes (Extended Data Table 5, see Methods). We then clustered differential expression profiles between healthy and regenerating or embolized conditions for genes with non-concordant zonation profiles, and identified groups of genes that similarly differed from healthy (Fig. 3i-l). In both PVE conditions, we observed that genes differing from healthy were associated with cellular respiration (E.g. CYB5A, NDUFA3 in regenerating; CP, MT-ND6 in embolized) and lipid metabolism (E.g. APOA5, CYP2A6 in regenerating; APOC3, APOC2 in embolized) (Extended Data Fig. 7c and 7d). In regenerating hepatocytes, we also observed genes such as GSTO1, SOD2, MT2A, and ASS1, which have been involved in response to toxic substances. These data show that metabolic homeostasis may be disrupted or in flux after the PVE procedure.

### Altered zonation patterns in sinusoidal endothelial cell states after PVE

We next examined zonation in liver sinusoidal endothelial cells (LSEC) after PVE (Fig. 4a). Our analysis in healthy fresh and frozen tissue datasets revealed periportal and pericentral populations consistent with previous reports^7,9,12^ (Extended Data Fig. 4). To identify zonation in PVE samples, we first annotated liver endothelial cell heterogeneity by subsetting and reclustering endothelial cells from the combined healthy and post-PVE datasets (Fig. 4b,c). This revealed 9 molecularly distinct LSEC subpopulations, as well as clusters of non-LSEC ECs, lymphatic ECs, and ECs in the G2M/S phase of the cell cycles (Fig. 4c,d). Based on marker gene expression, we cataloged the LSEC subpopulations as periportal (MGP, AQP1, EDNRB, CLEC14A), pericentral (CLEC1B, CLEC4G, CLEC4M, and FRZB), midzonal (periportal/pericentral markers), fenestrated (PLVAP, RBP7), remodeling (CTGF, IGFBP3, ANGPT2), interferon (CXCL10, IFI44L, ISG15, IFIT3), as well as three subpopulations characterized by the expression of mitochondria- and stress-associated genes, which may be a direct response to the shear stress associated with the changes induced by PVE.

**Fig. 4.**
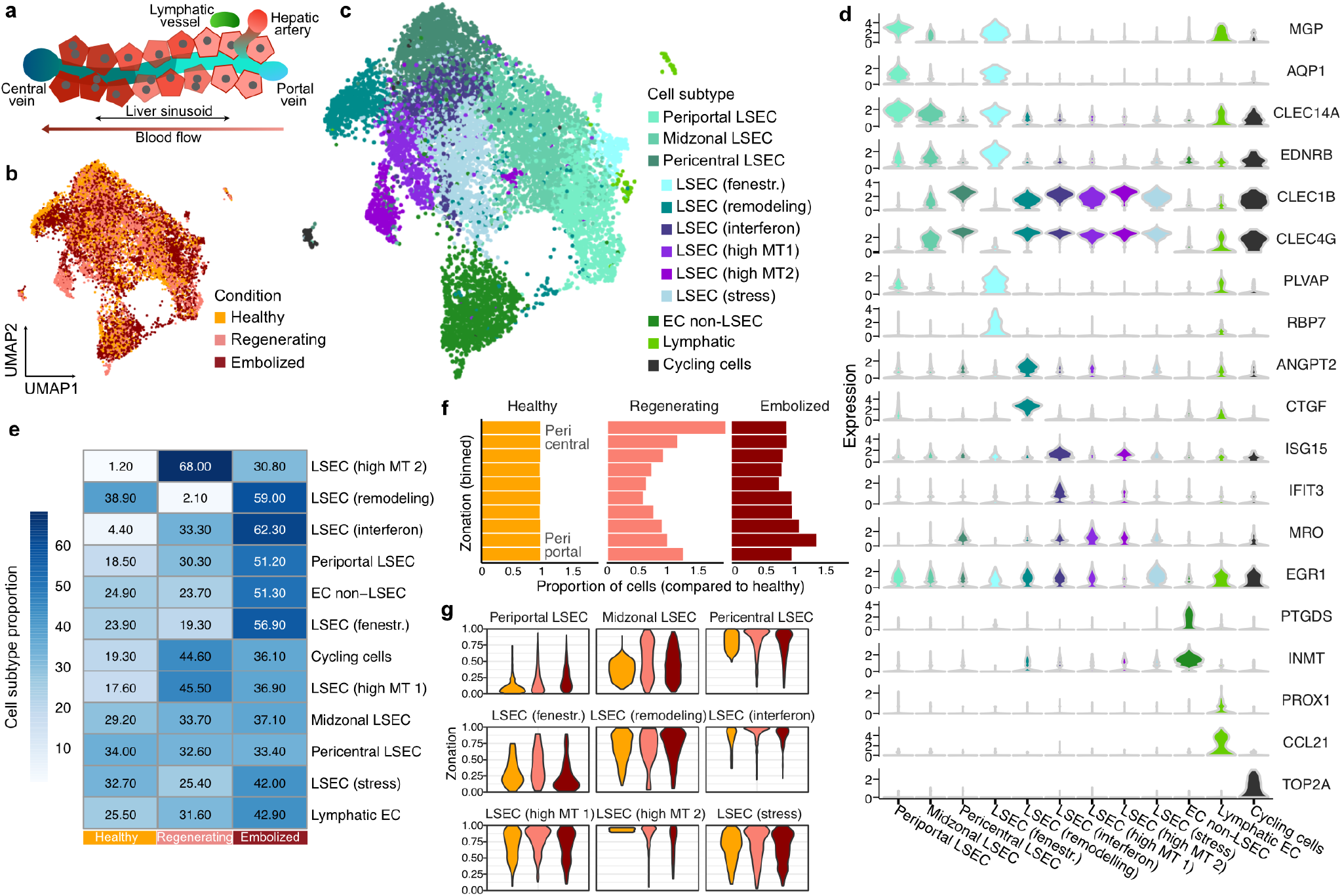
Liver endothelial cell heterogeneity and inferred zonation after portal vein embolization. a, Schematic illustrates diversity of endothelial cells along the portal-central axis within a liver lobule. b-c, UMAP plots representing combined ECs from all medical conditions are colored by condition (b) or annotated subpopulation (c) (EC, endothelial cells; LSEC, liver sinusoidal endothelial cells; fenestr., fenestrated; MT, mitochondrial). d, Violin plots show distribution of normalized marker gene expression for endothelial cells subpopulations. e, Proportion of each endothelial subpopulation across conditions (healthy, left; regenerating, middle; embolized, right). f, Histogram shows comparison of LSEC (periportal, midzonal, and pericentral LSEC) pseudozonation across conditions. LSECs from regenerating and embolized conditions were projected onto a reference healthy zonation trajectory. g, Distribution of each LSEC subpopulation from healthy, regenerating and embolized condition is shown along the pseudozonation trajectory.

We observed differential abundance of pericentral LSEC populations, with more cycling cells and high mitochondrial content cells in the regenerating condition, and an enrichment of interferon and ANGPT2+ remodeling cells in the embolized condition (Fig. 4e). Pseudo-zonation trajectory alignment to the healthy reference suggested a moderate shift towards pericentral identity in the regenerating liver, whereas LSEC from embolized samples had a slight inclination towards a periportal expression pattern (Fig. 4f, Extended Data Fig. 8a and 8b). We identified similarities in differential expression in regenerating and embolized LSEC populations in comparison to the healthy condition with a GO enrichment of differentially expressed genes in developmental, cell adhesion and migration programs (Extended Data Fig. 8c and 8d). Various genes also differed between the conditions (Extended Data Fig. 8e, Extended Data Table 6), yet these do not translate into any enriched GO Terms.

We projected each LSEC population to the healthy pseudozonation reference and found that fenestrated LSECs have a predominantly periportal signature, whereas remodeling, interferon-high, and stress LSECs primarily project pericentrally (Fig. 4g). Together with the divergent abundance of LSEC cell states in the three surveyed conditions, these inferred zonation differences shed light into the effects of PVE along the portal-central sinusoidal structure, with likely major effects on intercellular signaling within the liver lobules.

### The immune compartment has a pivotal role in modulating liver response to PVE

We next explored how PVE affects cell-cell communication in the liver (Fig. 5a). We first used the fresh tissue data to establish a finer-grained annotation of the various cell populations present in all three conditions with a particular focus on the immune compartment (Fig. 5b). We identified resident Kupffer cells as well as other macrophages, conventional and plasmacytoid dendritic cells (cDCs, pDCs), αβ- and γδ-T cells, B cells and plasmablasts consistent with previous studies^9,12^. We observed differential cell abundance of immune subtypes with macrophages being enriched in both post-PVE conditions and αβ-T and γδ-T cells being specifically enriched in the regenerating tissue (Fig. 5c). Cholangiocytes showed a lower abundance in both regenerating and embolized conditions, while hepatocytes showed a higher abundance post-PVE.

**Fig. 5.**
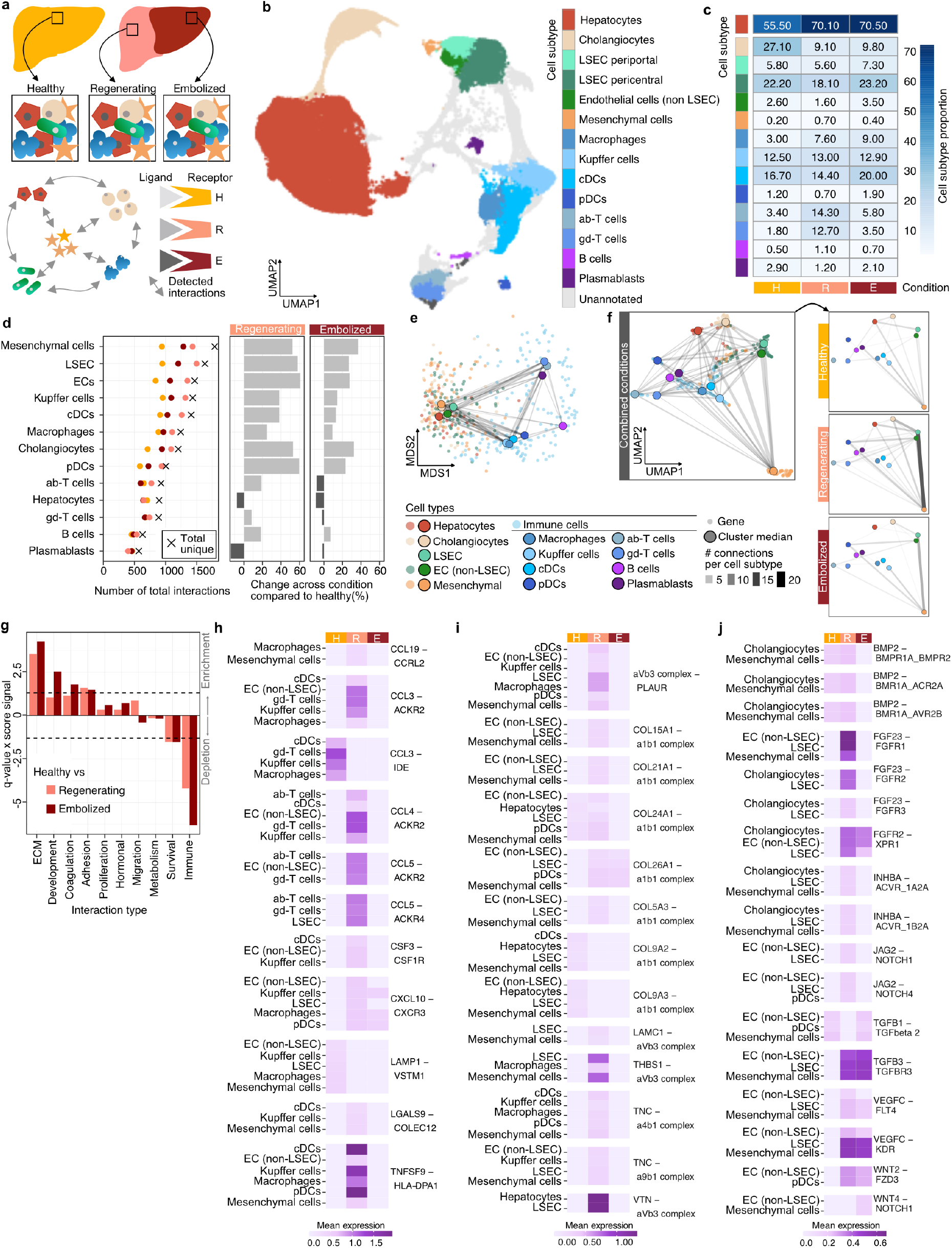
Intercellular signalling map during portal vein embolization reveals immune-cell coordination of response. a, Schematic illustrating ligand-receptor analysis across conditions. b, UMAP plot shows annotated liver cell subtypes (cDCs, conventional dendritic cells; pDCs, plasmacytoid dendritic cells; ab-T cells, alpha beta T cells; gd-T cells, gamma delta T cells). c, Heatmap shows subtype proportion across healthy, regenerating, and embolized conditions. d, Number of interactions involving each cell subtype per condition and in total (left dotplot), along with variation in the number of interactions between healthy and embolized (middle) or regenerating (right). e, Multidimensional scaling projection of ligand and receptor gene correlations, highlighting the role of immune-expressed receptors and ligands in signaling to non-immune cell types. f, UMAP plot of ligand and receptor genes (small circles, colored by cell type with highest average expression) with conditions combined (left) and for each condition individually (healthy, topright; regenerating, middleright; embolized, bottomright). g, Enriched (positive) or depleted (negative) interaction functions in variable interactions between healthy and embolized or regenerating. Top varying interactions involving h, immune i, adhesion and extracellular matrix components and j, development-related interactions in regenerating and embolized conditions compared to healthy.

Next, we aimed at understanding how the different cell populations together orchestrate the repair and regrowth of the hepatic tissue. We thus predicted cell-cell interaction events based on unique ligand-receptor pairings^30^ between cell subpopulations within each condition (Extended Data Table 7). Despite their low abundance overall, mesenchymal cells were the population with the most unique interactions, closely followed by LSECs (Fig. 5d left). Indeed, these two cell populations expressed a diverse array of extracellular ligands and transmembrane receptors, and displayed the largest amount of non-homotypic interactions (Extended Data Fig. 9). Most cell types increased the number of interactions in both post-PVE conditions compared to healthy, with the increase in interactions involving mesenchymal, endothelial, and myeloid cells (Fig. 5d). We observed that both αβ- and γδ-T cells followed a unique pattern of specifically increased interactions in the regenerating tissue (Fig. 5d). Curiously, hepatocytes showed a modest reduction of interactions in the post-PVE liver compared to healthy.

We summarized ligand-receptor pairs into a co-expression network using the gene expression correlations across cell types, and projected genes using MultiDimensional Scaling (MDS) (Fig. 5e). MDS components 1 and 2 clearly distinguished ligands and receptors expressed in immune cells from those found in other cell populations. Using a UMAP embedding to gain enhanced resolution into this interaction network, we found that upon PVE non-LSEC endothelial cells and mesenchymal cells establish new interactions with plasmacytoid Dendritic cells (pDC) and T cells (Fig. 5f). We performed gene set enriched analysis (GSEA) of the most variable interactions between conditions and found that interactions upregulated post-PVE were associated with cell adhesion, extracellular matrix (ECM) function, as well as development and coagulation, with the latter two being particularly strongly enriched in embolized tissue (Fig. 5g, Extended Data Fig. 10, see Methods). Conversely, both PVE conditions showed a depletion in survival and immune response-associated interactions.

Finally, we explored the particular ligand-receptor pairs that underly the differential interaction of cell populations in the different conditions. We found that upon PVE, T cells express various chemokines capable of influencing cell migration and morphogenesis (Fig. 5h), and pDCs appeared to have a role in coordinating cell adhesion and ECM structuring (Fig. 5i). It has been shown that the expression of ECM protein Tenascin-C (TNC) by mesenchymal cells in the regenerating liver may have an important role in modulating macrophage activation^31^, a mechanism important for the establishment of a pro-regenerative environment. We also observed an interplay between LSEC and hepatocytes via vitronectin (VTN) in regenerating samples, an interaction with likely regulatory implications in angiogenesis^32^. Interactions linked to developmental processes were represented in post-PVE liver (Extended Data Fig. 10), including PVE-specific upregulation of signaling through BMP, FGF, Wnt2, VEGF and TGF activation (Fig. 5j). For example, while TGFB3, a ligand previously reported to be crucial for liver development and regeneration^33,34^, was higher expressed in ECs, LSEC and mesenchymal cells in both PVE conditions, TGFB1 was not detected in the regenerating tissue. A possible reason could be the counteracting effects of BMP2 (present in healthy and regenerating samples), a molecule important in resolving liver fibrosis^35^. In addition, FGF family members were higher expressed in the regenerating cells, consistent with other tissue repair processes^36^. Also, in the TGF superfamily of proteins, INHBA signaling was solely captured in the regenerating tissue. This molecule was shown to have roles in tumor regeneration and pluripotency maintenance^37^. Altogether, these data revealed that diverse interactions were induced in the hypertrophic and atrophic conditions in response to PVE, and that therapeutic modulation of immune-dependent interactions may be harnessed to enhance desired outcomes.

## Discussion

Liver regeneration is a complex process frequently studied in animals, however it remains a major challenge to explore and understand how hepatic cells orchestrate liver replenishment in humans^18^. Animal models have provided information on the cellular sources and molecular pathways operating during liver regeneration^19^, including through the use of scRNA-seq^20,38–40^. Although this knowledge informs researchers and clinicians about regenerative processes likely occurring in humans, there is a lack of descriptive knowledge from primary human tissues that have undergone regeneration. These gaps emerge due to several reasons: obtaining regenerating liver tissue samples is usually not ethically feasible, available post-surgical human samples of regenerating liver are very scarce and difficult to obtain, and there are logistic and technical challenges associated with sample collection and time-dependent experiments as well as tissue processing and storage, respectively.

We generated a new resource to investigate human liver regeneration by establishing experimental protocols for surveying cell states in healthy fresh and frozen human liver samples, and showed the power and limitations of each approach. We established the first human liver regeneration transcriptomic atlas by sequencing more than 100,000 cells of healthy and post-PVE hepatic tissues. Our optimized protocols will allow access to frozen liver tissue banks and facilitate a more flexible investigation into regenerative processes, hepatic diseases, and malignancies. Second, we provided a detailed analysis of cell states, zonation patterns, and predicted intercellular interactions during two important physiological conditions, atrophy and hypertrophy. Our study covers different snapshots of regeneration because there is variation between individuals in dependence on their PVE duration. These snapshots include in varying shares early states of regeneration with focus on hepatocyte proliferation, and later states with focus on vascular growth and ductular reaction to reconnect the new hepatocyte mass to the circulation and drainage, respectively. We identified cellular expression profiles and pathways related to type I interferon response and cytokine signaling, as the general characteristics of the regenerating liver. This signaling is associated with the termination of hepatocyte proliferation^42^, indicating late regeneration stages of our samples. We also observed activation of developmental programs in cholangiocytes and endothelial cells, which are of crucial importance to re-establishing liver lobule structure and function. Our work will serve as a reference to test predictions to steer regenerative and developmental states using liver organoid model systems^43–45^.

PVE resulted in changes to the spatial organization and identity across hepatocytes and LSEC in the liver lobule. We predicted that hepatocyte zonation deteriorated in the embolized tissue, and hepatocytes had an augmented metabolic zonation profile. Post-hypertrophied hepatocyte zonation was partially restored in the regenerating samples, yet all post-PVE hepatocytes presented some metabolic alterations compared to the healthy liver, in particular at the level of lipid metabolism and energy production. Future studies using different post-PVE time points will be able to explore the long-term recovery of zonation profiles in the regenerated tissue, and focus on understanding regulation of these pathways to improve clinical recovery.

We found that LSEC populations retained zonation profiles in post-PVE tissues. However, we observed differences in cell state proportions, as well as stress, EC remodeling, interferon signaling, and cell cycle regulation as differential between conditions. These processes are reminiscent of previous work from mice, in which inductive angiocrine signals from sinusoidal endothelium stimulate hepatocyte proliferation, and the secreted ligand Angiopoietin 2 orchestrating phases of LSEC and hepatocyte proliferation^46,47^. Further work may unveil the location and role of these subpopulations in reorganizing the liver tissue and the sinusoidal vessels.

Beyond this, we uncovered a plethora of cell-cell interactions associated with liver regeneration. We identified a pivotal role for mesenchymal cells, as well as various immune cell populations. While Kupffer cells and other myeloid populations are expected players in liver regeneration^42,48^, we were able to further untangle contributions from pDCs, as well as αβ- and γδ-T cells. Most detected interactions were associated with ECM maintenance and cell adhesion, as well as signaling pathways related to liver development, such as WNT and TGFβ. Curiously, hepatocytes were one of the cell types least involved in cell-cell communication events in both PVE conditions. This highlights the role of non-parenchymal cells in orchestrating tissue morphogenesis after the initial wave of hepatocyte proliferation has occurred in PVE.

Altogether, understanding hepatic growth, regeneration, and degeneration is a topic of growing importance in the medical and bioengineering fields^43,49^. Our work delivers a human PVE atlas which can be used as a reference and blueprint for studying the mechanisms of human liver regeneration.

## Methods

### Experimental model

#### Human liver tissue samples

Human adult liver tissue samples were obtained from macroscopically “healthy” tissue that remained from resected human liver of patients with primary or secondary liver tumors or benign liver diseases with or without pretreatment with portal vein embolization. Informed consent of the patients for the use of tissue for research purposes was obtained according to the ethical guidelines of Leipzig University Hospital (006/17-ek, 21 March 2017, revised and renewed 12 February 2019). Acquired tissue from portal vein embolized livers included samples of regenerating (R) and embolized (E) tissue whereas tissue samples from benign liver diseases were defined as quiescent healthy controls (H) (Extended Data Table 1).

### Experiments

#### Isolation of human liver cells from fresh tissue

Isolation of the primary human hepatocytes (PHH) and non-parenchymal cells (NPC) from the liver tissue was performed as described previously^50^. Briefly, PHH and NPC were isolated from the same tissue sample simultaneously by a two-step EDTA/collagenase perfusion technique and then purified by Percoll density gradient centrifugation. To obtain different cell types from NPC fraction, this cell suspension underwent two different centrifugation steps: 300g for 5 min to get liver endothelial cells, mesenchymal cells and Kupffer cells; 650g for 7 min to get the majority of Kupffer cells. Finally, 3 cell suspensions, PHH, NPC300 and NPC650, were used to prepare a single-cell RNA-seq experiment. In case of samples with medical conditions, the regenerating and embolized tissues, the isolation procedure was made in the same manner.

#### Single-cell suspension preparation and single-cell RNA-seq experiment

Single-cell RNA-seq experiments were performed using a 10X Genomics platform. Before loading on a microfluidic chip, suspensions of PHH and both NPC fractions were washed and filtered at least twice in ice-cold 1X HBSS with-out Ca2+ and Mg2+ (HBSS w/o, Sigma) to remove tissue and cellular debris and to get individual cells in media that is compatible with the downstream experimental steps. Wide-bore pipette tips were used working with PHH to avoid premature cell lysis, while p1,000 and p200 pipettes were used gently resuspending NPC cell pellets.

Preparation steps of the single-cell suspension were made on ice and every time washed samples were spun down using a 4°C cooled centrifuge: PHH at 50g for 5 min, NPC300 at 300g for 5 min and NPC650 at 650g for 7 min. Finally, to generate single-cell suspension, PHH suspension was filtered through 40 and 30 and both NPC fractions through 30 µm diameter cell strainers. Cell viability and concentration were assessed using a cell analyzer (MuseTM Cell Analyzer, Luminex Corporation). NPC300 and NPC650 were pooled 1:1 and loaded on a 1 channel and PHH on a 2 channel of the Single Cell A and B chips targeting 6,000-8,000 cells per each sample. All steps of the single-cell suspension preparation for the regenerating and embolized tissue samples were executed following the healthy tissue protocol.

The next steps were conducted as described in the Chromium Single Cell 3’ Reagent v2 and v3 Kits. In brief, after generation of the droplets with the single cells and barcoded beads, cDNA synthesis was performed. Next, droplets were broken, cDNA was amplified and libraries were constructed with different Chromium i7 Sample indexes in order to record sample assignment during computational analysis. Finally, single cell libraries were run paired-end (28 bp, 8 bp, 100 bp) on an Illumina HiSeq2500 platform on 2 lanes. Experimental summary metrics can be found in Extended Data Table 8.

#### Frozen human liver tissue dissociation into single-nucleus suspension and flow cytometry sorting

Human frozen liver tissue samples were dissociated into single-nuclei combining liquid homogenization cell lysis with Dounce homogenizer and detergent-based lysis methods. All steps of the nuclei isolation were performed on ice with precooled solutions and using 4°C mode centrifugation. The dissociation protocol that was previously used on brain tissue^51^, was optimized here to maximize the nuclei isolation for the liver tissue. The protocol after the optimization included the following steps: first, thawing tissue sample was cut into smaller pieces, minced and transferred into a glass dounce homogenizer. 30 strokes of pestle A were used to homogenize the tissue in 0.3 M Sucrose (Sigma) solution including 0.002 M EDTA (Thermo Scientific), 1% BSA (Serva) and 1% Tergitol solution (Sigma). After 5 min of incubation, next 30 strokes of pestle B were applied to finalize the disruption process and deliberate nuclei from the cells into suspension. Homogenized solution was centrifuged at 600g for 5 min and the nuclei pellet was washed twice in PBS solution (0.002M EDTA, 1% BSA, 0.2 U/ul RNase Inhibitor (Thermo Scientific)). Finally, to remove any aggregate and debris the nucleus suspension was filtered through a 30 µm diameter strainer and resuspended in PBS solution (1% BSA, 0.2 U/ul RNase Inhibitor). Further, to enrich for individual nuclei, the suspension was sorted by applying a 4-way purity mode based on the selected DAPI positive nuclei population (1:1000, BD Pharmingen) using forward and side scatter gating strategy (FACS). These nuclei were sorted in bulk and kept on ice for >30 min. To ensure that the sorted nuclei were intact, they were stained with DAPI (1:500) to inspect under the fluorescence microscope and finally counted using a hemocytometer before a single-nucleus RNA-seq experiment.

#### Single-nucleus RNA-seq experiment

Sorted single-nucleus suspensions were loaded on a Single Cell B chip to generate single-nucleus gel beads in emulsion on a 10X Chromium controller. Single-nucleus RNA-seq libraries were prepared following the protocol of the Single Cell 3’ Reagent Kit v3 and sequenced paired-end (28 bp, 8 bp, 100 bp) using an Illumina HiSeq2500 platform.

#### Cross-sectional Imaging

For liver segmentation and volume measurement 3D-datasets of both computed tomography (CT) and magnetic resonance imaging (MRI) were used.

CT was acquired as a 128-slice multidetector helical intravenously contrast enhanced (100-120 mL iomeprol; Imeron 400, Bracco, Milan, Italy) scanner (Ingenuity, Philips, Best, The Netherlands). Contrast phase scanning was adjusted to the necessity of visualization of all liver architecture relevant for segmentation (i.e., liver veins, portal vein branches (scan delay of 70-90 seconds after injection). Primarily axial reconstruction of images in 1-2.5 mm slice thickness (increment, 1).

MRI (1.5 T, Magnetom Aera, Siemens, Erlangen, Germany) was acquired with the use of 0,1 mL/kg body weight gadoxetic acid (Primovist/Eovist, Bayer, Leverkusen, Germany), in the early dynamic phases and negative contrast of blood vessels in the hepatobiliary phase (15-25 min. delay after injection) and a fat saturated breath hold T1-weighted interpolated 3D-sequence (VIBE).

All image data sets were digitally archived (DICOM format), pseudoanonymized, and exported to a dedicated postprocessing workstation (Lenovo ThinkStation, Lenovo, Beijing, China) inside the institutional network.

Volume segmentation was done semi-automatically by outlining the liver surface excluding large hilar vessels and interceptions with a dedicated post-processing tool (3D Slicer, open source software). Virtual resection along anatomic landmarks were performed to assess the future liver remnant before and after the portal vein embolization regarding the outcome parameters volume, vessel architecture and tumor progression. 3D-Visualization of the 3D model was performed using the Blender software (Blender Foundation, Amsterdam, Netherlands).

#### H/E staining and Immunohistochemistry

For investigation of tissue sections human liver tissue samples (n=3 per condition, Extended Data Fig. 6) were fixed with paraformaldehyde (PFA, Carl Roth, Karlsruhe, Germany), embedded in paraffin, sectioned in to 3.5 µm thick slices using a microtome (MicromHM430, Thermo Fisher Scientific, Waltham, MA, USA), and mounted on slides.

For Hematoxylin and Eosin (HE) staining the tissue sections were rehydrated. Then the tissue sections were incubated in Hämalaun (Merck, Darmstadt, Germany) for nuclei staining for 5 min at RT and washed under rinsing water for 10 min. Afterwards the tissue sections were stained with Eosin G-solution (Merck, Darmstadt, Germany) for 3 min. Finally, the tissue sections were dehydrated and the slides were embedded in the non-aqueous mounting medium Entellan (Merck, Darmstadt, Germany).

The immunostaining of tissue sections and their microscopic evaluation was performed as described previously^52^ with the following modifications. Briefly, tissue sections were rehydrated and epitopes were retrieved. Then, tissue slices were blocked for endogenous peroxidase activities and for unspecific binding. For detection of GLUL, HAL, SAA1/2, IGFBP2, CYP3A4 specific primary antibodies (all Abcam, United Kingdom, Cambridge) were used (Table 9). Antibodies were diluted in TBS (Sigma, Munich, Germany) with 1% BSA (Sigma, Munich, Germany) and 0.03% TritonX-100 (Sigma, Munich, Germany).

The antibodies against the targets were visualized using Peroxidase-conjugated secondary antibodies (Table 9). Secondary antibodies were diluted in TBST (TBS + 0.5% Tween20 (Sigma, Munich, Germany)). Detection was performed by using 3,30-Diaminobenzidine (DAB, Sigma, Munich, Germany).

The EnVision+Dual Link System-HRP (Dako, Glostup, Denmark) was used according to manufacturer instructions when the targets showed low expression in the tissue sections. All reactions were stopped and cell nuclei were stained with hematoxylin. Finally, the slices were dehydrated and embedded using Entellan (Merck, Darmstadt, Germany).

For immunofluorescence (IF) staining the above described procedure was repeated without blocking for peroxidases. For visualization the secondary antibodies ALEXA Fluor 647 donkey anti rabbit was used and diluted as described above. Cell nuclei were stained with Hoechst. Finally, the slides were embedded in Mowiol-488 (Carl Roth, Karlsruhe, Germany). Negative controls for DAB and IF stainings were made from all donors and treated in the same way but without usage of a primary antibody.

#### Imaging analysis

Whole slide images of IF stainings were captured in fluorescence mode using a Slide Scanner (AxioScan Z1, Carl Zeiss, Oberkochen, Germany) with a 20/0.8 M27 Plan-Apochromat objective with channel 1 (Target) light source 630 nm, light source intensity 50% with extinction wavelength 631 nm and emission wavelength 647 nm and with channel 2 (Hoechst) fluorescence light source 385 nm, light source intensity 14,53% with extinction wavelength 353 nm and emission wavelength 465 nm with an Axiocam 506m as imaging device. Resulting images were stored as raw data in the Carl Zeiss proprietary image pyramid format (CZI) with an object-related nominal pixel size of 0.227 µm x 0.227 µm.

Whole slide images of DAB and H/E stainings were captured in transmitted light mode using a Slide Scanner (Pannoramic Scan 2, 3DHISTECH, Budapest, Hungary) with a 20/1.6 Plan-Apochromat objective and stored as raw data in the MIRAX Virtual Slide Format (MRXS) with an object-related nominal pixel size of 0.243 µm x 0.243 µm.

Tissue sections were visually investigated for lobular architecture (n=3 per condition, analog to Extended Data Fig. 6). For that we used sequenced sections from immunofluorescence stainings for GLUL and Hoechst (pericentral hepatocytes and cell nuclei) and HAL and Hoechst (periportal hepatocytes and cell nuclei) to identify the lobular vessel architecture and lobular borders. For the further imaging analysis, the Hoechst channel from the GLUL staining was used. Here, feature regions containing identifiable liver lobules were stored from Zeiss ZEN in 10000 × 10000 Pixel Tiles as Portable Network Graphic (PNG). Clearly identifiable lobules were donor and condition dependent resulting in n = 15 lobules for healthy, n = 12 lobules for regenerated and n = 10 lobules for embolized tissue, respectively. Fiji (ImageJ) was used to preprocess the selected feature regions of the tissue. In a first step the coordinates of the lobule boundaries and the location of the central vein were set by hand and stored as “Comma Separated Values” (CSV) files for later imaging processing. Between each coordinate of the lobule boundaries and its corresponding central vein a portal-central axis results. The number of axes was at least six but finally dependent on lobule shape complexity and was tried to keep as close to six as achievable.Then a gaussian filtered version (sigma=20) of the image was subtracted from the original to correct staining artefacts and normalize overexposed regions and the background. After that the image was stored as “Tagged Image File Format” (TIFF). The density investigation started with an approximate estimate of the nuclei centers and measurement of their spatial locations. For that an Otsu threshold was used to filter the detected nuclei and reduce the amount of nonspecific signals resulting in a binarized image. The feature region of the nuclei in the binarized image were processed with morphological filters to remove small white noise and fill holes. For that we use morphological opening and closing. Regions near to the center of a feature region are defined as sure foreground and regions far away to a feature region are defined as background. The distance transformation provides the pixels known as certain of a region that belongs to a nucleus. With a weighted threshold of the maximum distance transformation value we received sure foreground regions of an identified nuclei indicated as a marker. A connected component analysis labels these regions with any positive integers to separate these from the background. Processing these markers with a marker-based watershed method allows to allocate the unclassified regions of the image belonging to nuclei regions or not. The resulting region property provides sure coordinates of spatial locations of nuclei.

To get the spatial dependent location density we use the most popular spatial analysis technique called kernel density estimation (KDE). KDE is a statistical method and a nonparametric way to estimate the probability density function of a random feature and correlates it to features in its neighborhood. By correlating each feature with a kernel function and summing up all the weighted overlapping regions of the kernel we get the probability density as a level of spatial distribution of nuclei density. The kernel is chosen as a multivariate gauss function with a fixed bandwidth of 0.2 for all images to ensure comparability of lobules from different sections among each other. The bandwith was empirically determined by analyzing 3 of the control samples by using the scott algorithm for bandwidth estimation.

The resulting spatial density distribution of nuclei was plotted as a heatmap. In a next step we transferred the lobule coordinates into the heatmap and extracted the density values along the earlier defined portal-central axis of a lobule (n = 121 axes for healthy, n = 84 axes for regenerated tissue and n = 76 axes for embolized tissue). The portal-central density distribution of a condition was plotted as the mean density values along the portal-central axes including standard deviation.

### Single-cell RNA-seq computational analysis

#### Data processing after sequencing (Cell Ranger pipeline)

We used the Cell Ranger software (https://support.10xgenomics.com/single-cell-gene-expression/software/overview/welcome) to process the sequenced RNA libraries and generate gene expression count matrices for the analysis. We first transformed Illumina intensities, raw base call (BCL) files into reads using cellranger mkfastq. Next, we ran cellranger count to align the reads to the human reference genome (GRCh38) using RNA-seq aligner STAR with default parameters (Extended Data Table 10). Uniquely mapped reads were based on barcodes and unique molecular identifiers (UMIs) assigned to cells and genes (ENSEMBL release 84) respectively. Read counts for a given gene and cell that are represented by a Chromium cellular barcode and UMI were used as an input for the subsequent expression analyses.

#### Single-cell data filtering and normalization

Prior to any processing, scrublet^53^ was used to assign a doublet score to all cells in each fresh tissue dataset. We used the R package Seurat (version 3.0)^54^ to process gene expression count matrices. We first applied SCTransform to normalize molecular counts, scale and identify variable genes^24^ within each dataset separately. Cells in each sample were then finely clustered (Louvain algorithm, resolution = 10) and the average doublet score was calculated to identify small groups of similar doublets. Quality control filtering was done by applying the thresholds outlined in Extended Data Table 11.

#### Integration of single-cell data

Samples were analyzed in two groups: healthy only and all conditions (healthy, regenerating and embolized). Both groups were integrated using CSS^23^. For the healthy data, integration was done using all common genes and the first 30 principal components. For integration of all samples, the top 3,000 variable genes from the healthy, regenerating and embolized samples, as well as the top 100 marker genes from each cell type identified in the healthy dataset were selected. These were used to do a PCA on the full data, of which the top 50 principal components were used. The genes considered allowed a coverage of the biological variability in all conditions and present populations, despite the unbalanced representation of cell types in each sequenced fraction (Hepatocytes and Non-parenchymal cells).

#### Dimensionality reduction, clustering and annotation of single-cell data

Projection with UMAP^55^ and clustering, both for the healthy and for all combined datasets, was performed using all dimensions obtained from CSS.

For the healthy data, clusters were obtained using Louvain clustering with 0.9 resolution, and markers were detected for these populations using Seurat’s FindAllMarkers function (pseudocount.use=0.1, logfc.threshold=0.2, adjusted p-value<=0.05). Some clusters (9, 12, 19) were further individually subclustered to identify specific endothelial, T cell, and pDC/B cell populations, respectively. Annotation was done based on the general and subclustering identified markers, which resulted in some smaller clusters being merged under the same label. Cells were also grouped into five “major cell types” (Fig. 1b), and their marker genes were also calculated using Seurat’s FindAllMarkers function (pseudocount.use=0.1, logfc.threshold=0.2, adjusted p-value<=0.05).

Clustering of combined datasets used the Louvain algorithm with 1.1 resolution. Markers were detected for the identified clusters using Seurat’s FindAllMarkers function (pseudocount.use=0.1, logfc.threshold=0.2, adjusted p-value<=0.05). Clusters 5, and 19, as well as clusters 6, 22, 26, were subclustered to identify more specific types of endothelial cells, macrophages, and T cells, respectively. All clusters and subclusters were annotated using their top marker genes, and resulted in some clusters being merged into the same cell population.

#### Identification of LSEC subtypes in the combined single-cell data

Subsets for LSEC were identified by reclustering previously annotated ECs in the combined dataset, followed by data renormalization. These cells were then filtered for non-endothelial and doublet populations based on previously reported LSEC marker genes^6,9^ and cells expressing genes from other cell types, respectively. Marker genes for each remaining cluster after filtering were determined using Seurat’s FindAllMarkers function (pseudocount.use=x, logfc.threshold=x, adjusted p-value<=0.05). This methodology allowed for the identification of bona fide LSEC cells (periportal, midzonal, and pericentral), as well as LSEC populations with unique expression profiles, cycling endothelial cells, lymphatic EC and other non-LSEC EC likely originating from the portal or central veins (Fig. 3).

#### Zonation signature creation in healthy, regenerating and embolized tissue hepatocytes

Information on expression of previously established human and mouse zonation marker genes^3,12^ was used to identify portal- and central-zone hepatocytes within healthy and post-PVE samples. For each of the three conditions we generated a combined zonation expression signature based on portal and central expression markers. For each gene in both gene sets we calculated the z-normalized expression value across all cells. We then transformed the resulting expression values into the range of between 0 and 1 by subtracting the min expression and dividing by the maximum expression per gene across cells. For each zone-specific gene set we calculated the sum of the normalized gene expression values in a given cell. Per cell we generated combined expression signatures by adding the negative portal signatures to the central signatures. Based on these scores clusters in the UMAP were defined to be showing portal or central specific expression signatures.

For this analysis healthy, regenerating and embolized tissue hepatocytes were subset individually from the UMAP of combined datasets based on previously annotated cell type markers. Within each dataset, cells were renormalized using the SCTransform function in Seurat. Four PCs were then used to project cells in UMAP space.

#### Comparing healthy, regenerating and embolized liver samples

For each annotated cell type, Seurat’s FindMarkers function was used to obtain the differentially expressed (DE) genes between pairs of conditions (Fig. 2f). A maximum of 10,000 cells was used for each condition. Additionally, in order to account for the sequencing depth, prior to calculating the DE genes for hepatocytes, the seqgendiff package^56^ was used to downsample the UMI counts for all conditions, taking the minimum median of UMI counts of the three conditions as reference (Extended Data Fig. 5b). Genes encoded in the Y chromosome were disregarded - male donors were only present in the regenerating and embolized conditions, as well as DE genes between conditions that have been detected as marker genes for other cell types that likely appeared due to ambient RNA contamination.

#### GO Term enrichment analysis

Enrichment for GO Terms was performed using the function enrichGO from the clusterProfiler^57^ package, using the all terms in the org.Hs.eg.db package database, with a q-value cutoff of 0.05. The genes considered for analysis were previously identified as DE with an adjusted p-value <=0.05 and a logFC>0.3. All genes tested for differential expression were used as a background set for the analysis. For plotting (Fig. 2k-n), significant GO Terms for each condition were clustered based on their gene similarities using hierarchical clustering, and then grouped into 6 clusters. The term with the lowest p-value per cluster was chosen as a representative to be plotted.

#### Identification and comparison of hepatocyte and LSEC zonation across medical conditions

A steady-state transcriptomic zonation reference was established by identifying a latent axis ordering healthy hepatocytes (Fig. 3) and LSEC (Fig. 4) independently, using DiffusionMaps from the destiny package^58^.

For hepatocytes, contaminating non-hepatocytes were first filtered, followed by renormalization and PCA. The first 4 PCs were used as input for DiffusionMaps, and the ranked DC1 dimension, normalized to values between 0 and 1, was defined as the healthy hepatocyte pseudozonation trajectory. Genes varying along this trajectory were determined by parametric ANOVA on a Generalized Additive Model meant to predict gene expression dependent on the pseudozonation trajectory, modelled as a natural spline with 3 degrees of freedom.

LSEC zonation was determined by applying DiffusionMaps to the first 10 PCs of bona fide LSEC populations - Periportal, Midzonal, and Pericentral LSEC (Fig. 4). This first step identified a few outlying cells. These were removed, and the remaining data was renormalized and projected with PCA. DiffusionMaps was run on the first 10 PCs, and the identified DPT was used as a pseudozonation trajectory, after ranking and normalization to the 0-1 interval. Genes varying along this trajectory were determined similarly to those for hepatocytes.

Comparison of hepatocyte and LSEC zonation in the regenerating and embolized liver to the established healthy reference was done by selecting the top 1,000 varying genes in the healthy pseudozonation, and used them to train a generalized additive model to predict the pseudozonation variable (Extended Data Fig. 7a and 8a). This model used a beta distribution for error modelling with a logistic link function, which guaranteed that the predicted trajectory would be in the interpretable 0-1 range. In each condition, varying genes were determined as described above.

Genes varying in the hepatocytes were determined as differing between pairs of conditions if their Spearman’s rank correlation coefficient was lower than 0.3. The fitted expression of these genes was clustered using Euclidean distance and the ward.D2 method for healthy vs regenerating and healthy vs embolized, to identify groups of genes differing in similar ways (Fig. 3i-l).

LSEC varying genes were compared between conditions using Spearman’s rank correlation on the fitted values. A correlation coefficient greater than 0.3 indicated a similar behavior between conditions, whereas values below that were considered a different or opposite behavior between conditions. To illustrate this, Extended Data Fig. 8e shows the top 30 similar genes of healthy vs regenerating and healthy vs embolized (PCC>=0.3), as well as the top 35 different genes (PCC<0.3) of the same comparisons; each group was obtained by clustered using Euclidean distance and ward.D2 method.

#### Identifying cell-cell communication events in healthy, regenerating and embolized liver

Ligand-receptor pairs mediating cell-cell communication events were detected within each condition using CellPhoneDB (version 2.0)^30^, based on the annotated cell types from the complete data integration (Fig. 5b). The detected ligands and receptors were then used to create two types of projections summarizing cell-cell communication in the healthy and post-PVE liver. We projected a graph showing all correlations greater than 0.3 between all ligands and receptors using multidimensional scaling (Fig. 5e), and summarized in the same coordinate space each cell type as the median coordinates of the ligands and receptors that are expressed in it at the highest level. We also projected the mean expression per cell type and condition of all ligands and receptors using UMAP (Fig. 5f), and identified the interactions that are unique to healthy or both PVE conditions.

#### Detecting enriched types of variable interactions per condition

For each interaction, in each condition, we obtained a vector encoding whether an interaction was detected in a given pair of cell types. We used these vectors to calculate the mutual information between healthy and regenerating and healthy and embolized samples for each interaction. The resulting values were then used for Gene Set Enrichment Analysis^59^ to determine enriched or depleted types of interactions (Fig. 5g). Interaction types were manually annotated based on literature searches, and can be found in Extended Data Table 7.

### Single-nucleus RNA-seq computational analysis

#### Single-nucleus RNA-seq data filtering, normalization and clustering analysis

We used the R package Seurat (version 3.0)^54^ to process gene expression count matrices. We first applied SCTransform to normalize molecular counts, scale and identify variable genes^24^. After manual inspection we applied per sample minimum and maximum thresholds on the number of detected genes in a given nucleus to exclude both nuclei with low RNA content and potential doublets (sn_H1: >150 and <1,700 detected genes; sn_H3: >200 and <2,000 genes; sn_H4: >70 and <1,100). In addition, we excluded nuclei with more than 10% of UMIs aligning to mitochondrial genes. The number of nuclei used in the analysis for each condition is provided in Extended Data Table 12.

#### Integration of the single-nuclei datasets using batch effect correction

Sample-specific preprocessed datasets were merged based on the 3,000 most variable genes Pearson residuals.

#### Cell type identification analysis in the merged single-nuclei healthy datasets

UMAP was used to represent the similarity of gene expression profiles between nuclei in 2D. The clustering of the healthy nuclei data revealed 5 major clusters. Clusters were assigned to cell types based on the presence of cell type marker genes that showed a significantly higher expression in a given cluster. DE was performed using Wilcoxon Rank Sum test between each cluster and remaining clusters.

#### Identification of zonation within hepatocytes and LSEC of fresh and frozen healthy liver tissues

We used the expression of previously established human and mouse zone-specific marker genes^3,9,12^ to identify portal and central hepatocytes (Extended Data Fig. 3, b-c) as well as periportal and pericentral LSEC (Extended Data Fig. 4, a-b) in both fresh and frozen tissue datasets. Portal and central expression signatures were calculated separately across these marker genes in each of the two cell types and datasets. Z-normalization was done per gene and the portal and central scores represented the sum across normalized portal or central marker gene expressions in a given nucleus or cell. The signature was then shown on UMAP embeddings of each cell type and processing protocol. For the frozen tissue dataset, a UMAP embedding containing previously annotated cell types was used. For the fresh tissue dataset hepatocytes and endothelial cells were projected using the top 4 or 15 principal components, respectively. Portal and central signatures were then used to define portal and central groups of cells. DE genes were identified between these sub-clusters in fresh and frozen hepatocytes and LSEC, respectively, by using FindMarkers function in Seurat with logFC threshold being at 0 and other default parameters. Fold changes between both zonation-linked DE analyses were significantly correlated for hepatocytes (Spearman’s rho = 0.19, p = 2.7×10-21) (Extended Data Fig. 3d) and LSEC (Spearman’s rho = 0.2, p = 4.6×10-27) (Extended Data Fig. 4c), suggesting that the tested sub-clusters between cells and nuclei shared a portal and central pattern of zonation.

For the hepatocytes the following genes HAMP, CRP, SDS, NAMPT, HAL, ID1 were identified as being higher expressed and were shared between both datasets in portal sub-clusters (logFC > 0.4 for fresh and logFC > 1 for frozen) (Extended Data Fig. 3d,e). Conversely, CYP2E1, CYP3A4, IGFBP1, BAAT, SLCO1B3, KLF6 showed higher expression in central sub-clusters of both datasets (logFC < -0.4 for fresh and logFC < -1 for frozen) (Extended Data Fig. 3d,e). For the LSEC, VIM, EMP1, SPRY1, LMNA showed particularly high expression in peri-portal subclusters (with the logFC > 0.9 for fresh and logFC > 2 for frozen) (Extended Data Fig. 4c,d) and CLEC4G, STAB1, CLEC4M, OIT3, CTSD, LYVE1, CTSL, CD14 were significantly higher expressed in peri-central clusters (Extended Data Fig. 4c,d) in both fresh and frozen tissue datasets.

#### Zonation-specific protein expression validation using Human Protein Atlas

Healthy liver immunostaining images (Extended Data Fig. 3f) were downloaded from the Human Protein Atlas^60^ for expression information analysis of genes showing portal and central specific expression in our dataset.

## AUTHOR CONTRIBUTIONS

AB generated the single-cell and single-nucleus data. TG, AB, ZH and MD analyzed the data. CK collected tissue samples and performed liver cell isolations. AB and TS performed single-nucleus suspension sorting using FACS with assistance from MS. JCE performed immunohistochemistry. RH performed imaging analysis. DS was responsible for liver surgery, patient acquisition, clinical data and CT/MRT analysis. TD was responsible for the PVE, CT/MRT imaging. MB and JH provided intellectual guidance for data analysis. AB, TG, MB, JH, GD, BT, and JGC designed the study. AB, TG, GD, BT, and JGC wrote the manuscript with input from all other authors. All authors read and approved the manuscript.

## ACKNOWLEDGEMENTS

We thank the Camp and Treutlein labs for helpful discussions, and S. Pääbo, K. Köhler, B. Nickel, B. Schellbach, A. Weihmann, J. Kelso of Max Planck Institute for Evolutionary Anthropology for supporting this project. We also would like to acknowledge Markus Morawski from Paul Flechsig Institute of Brain Research at University Leipzig for sharing the slide scanner AxioScan Z1 (Carl Zeiss) and Karsten Winter from the Institute of Anatomy at University Leipzig for sharing the slide scanner Pannoramic Scan 2 (3DHISTECH). JGC and BT are supported by grant number CZF2019-002440 from the Chan Zuckerberg Initiative DAF, an advised fund of the Silicon Valley Community Foundation. JGC is supported by the European Research Council (Anthropoid-803441) and the Swiss National Science Foundation (Project Grant-310030_84795). BT is supported by the European Research Council (Organomics-758877, Braintime-874606), the Swiss National Science Foundation (Project Grant-310030_192604), and the National Center of Competence in Research Molecular Systems Engineering. TG is supported by an EMBO Long-Term Fellowship (ALTF 738-2019). MD is supported by the European Union through Horizon 2020 Research and Innovation Program under Grant No. 810645 and the European Union through the European Regional Development Fund Project No. MOBEC008. JH and DS are supported by the German Research Foundation (DFG, Germany) grant number: HA3091/14-1, HA3091/12-1 and SE 1694/4-1, SE 1694/5-1, respectively.

## DATA AVAILABILITY

Raw and processed scRNA-seq and snRNA-seq data generated and used in this study have been deposited in ArrayExpress under accession XXXXXXX and Mendeley (http://dx.doi.org/10.17632/yp3txzw64c.1), respectively. Imaging data used in this study for lobular density investigation was deposited in Zenodo.org doi: 10.5281/zenodo.4772378.

## CODE AVAILABILITY

R notebooks and scripts used in this analysis can be found in https://github.com/tomasgomes/liver_regen and https://github.com/ReneHaensel/Liver_regen_cell_Atlas

