## Extended Data Figures for "Cell atlas of the regenerating human liver after portal vein embolization"

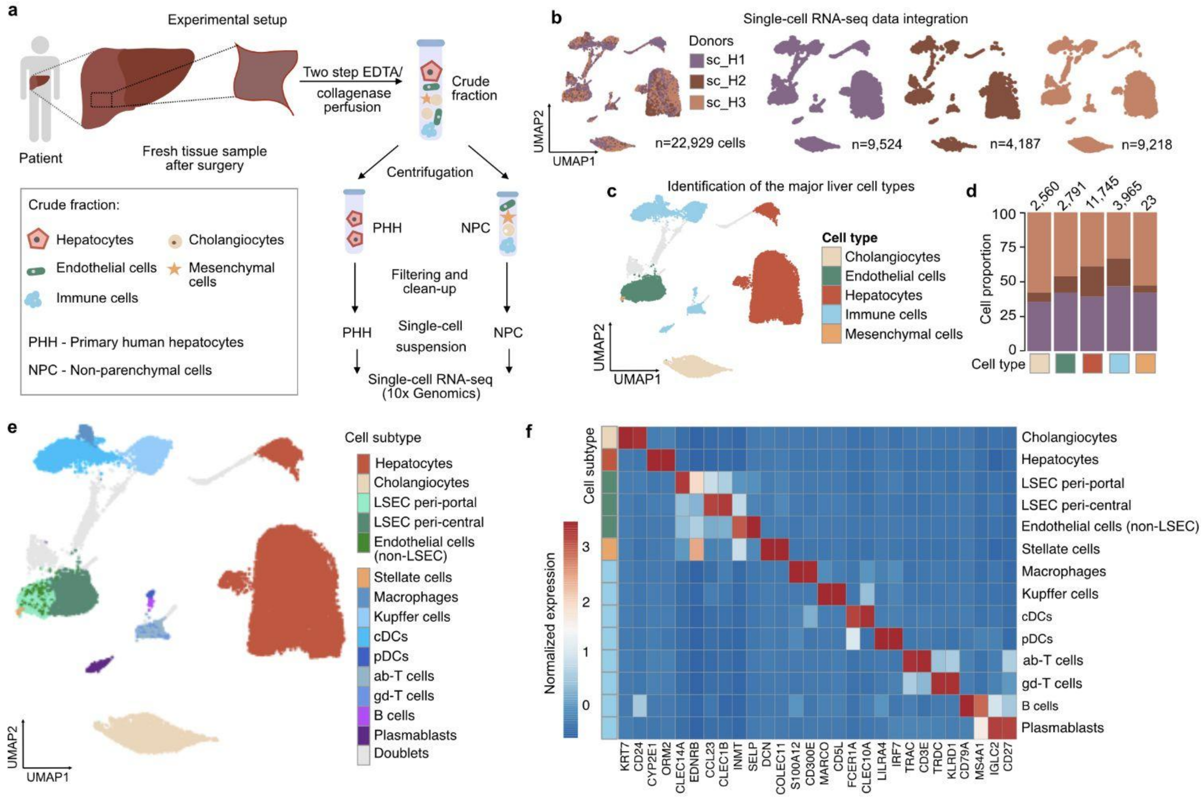


**Extended Data Fig. 1 | Single-cell RNA-seq from fresh human liver control specimens reveals different cellular subtypes**. **a**, Experimental schematic illustrating isolation of the primary human hepatocytes (PHH) and non-parenchymal cells (NPC) from patient-derived tissue samples (n=3). Two cellular fractions were processed and used separately to perform sc RNA-seq. **b, c**, UMAP plots of scRNA-seq data from three fresh healthy (H) liver donors, colored by individual donor contributions (**b**), and by the identified major cell types (**c**). **d**, Barplots show proportions of cells from three donors across cell types, along with the absolute number of cells detected per cell type (top). **e**, UMAP showing different cell subtypes within the liver. **f**, Heatmap showing expression of cell marker genes per annotated cell subtype (genes are represented in columns, cell subtypes in rows).

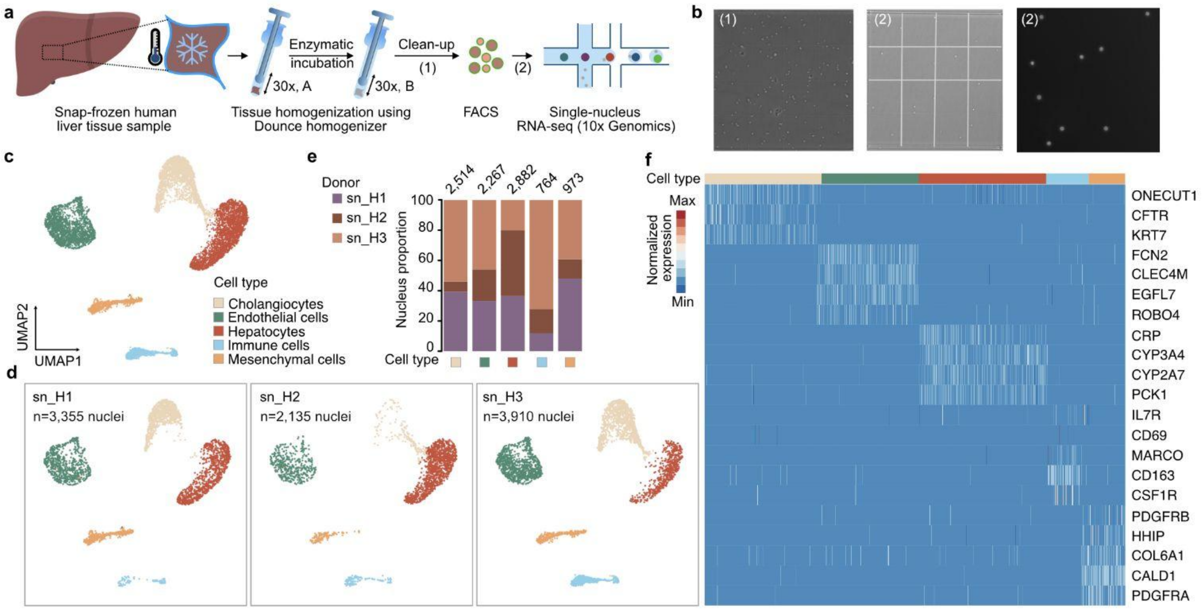


**Extended Data Fig. 2 | Single-nucleus RNA-seq from snap-frozen healthy human liver specimens. a,** Workflow illustrating experimental steps of single nucleus isolation from snap-frozen human liver tissue. **b,** Phase-contrast micrographs show representative single-nucleus suspension stained with DAPI after filtration and clean-up before (1) and (2) after FACS (10x magnification). **c, d**, UMAP plots of single-nucleus transcriptomes of three frozen healthy liver samples colored by cell type and showing merged (**c**) or individual (**d**) contributions of each donor. **e**, Barplots showing proportions of cells from three donors across identified cell types are shown together with the absolute number of cells detected per cell type (on top). **f**, Heatmap shows normalized expression of cell marker genes per annotated cell type (nuclei are represented in columns, genes in rows).

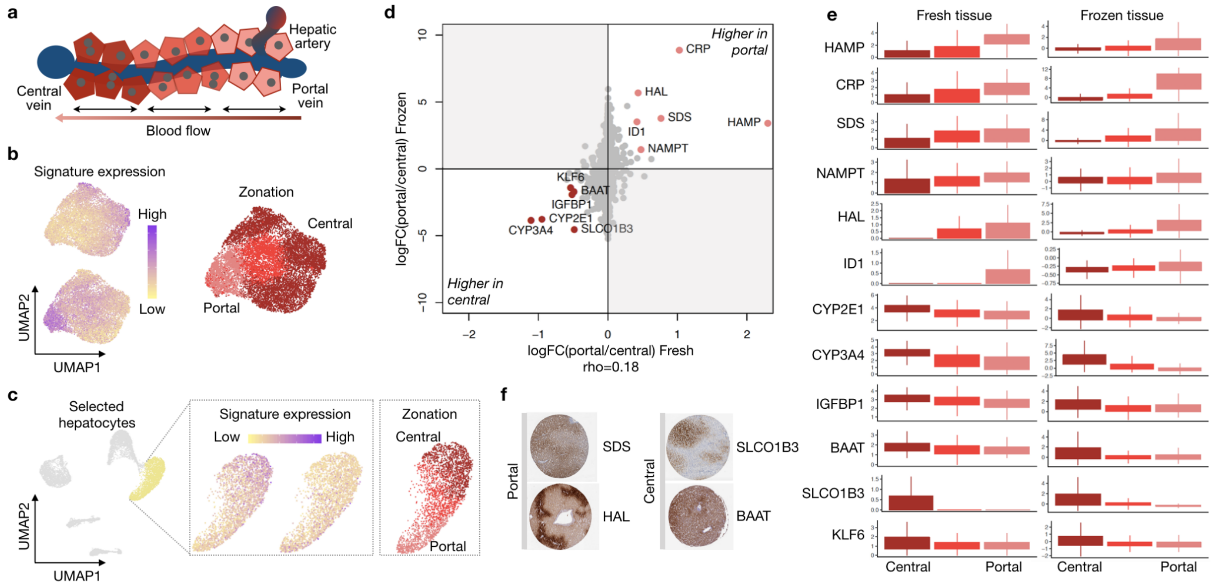


**Extended Data Fig. 3 | Identification of the spatial pattern of zonation in healthy tissue hepatocytes.** **a**, Schematic of portal-central axis of zonation in hepatocytes within liver lobule. **b**, UMAP plot of fresh tissue hepatocytes showing average gene expression of portal (bottom left) and central marker genes (upper left) and annotated subclusters (right) based on zonation signature expression (see Methods). **c**, UMAP plot of frozen tissue with hepatocytes showing average gene expression of portal (bottom left) and central marker genes (upper left) and annotated subclusters (right) based on the zonation signature expression (see Methods). **d**, Fold change showing differential expression in hepatocytes from portal and central subclusters in fresh (x-axis) and frozen (y-axis) tissue data (Spearman's rho=0.19, p=$2.7\times10^{-21}$). Genes sharing the zonation pattern, i.e. genes with abs(logFC)>0.4 for fresh and abs(logFC)>1 for frozen are shown in light red (portal) and dark red (central), respectively. **e**, Expression of identified genes across hepatocytes from portal and central subclusters in fresh and frozen tissue datasets. **f**, Immunostaining (Human Protein Atlas) of SDS, HAL and SLCO1B3, BAAT proteins representing portal and central zonation pattern, respectively.

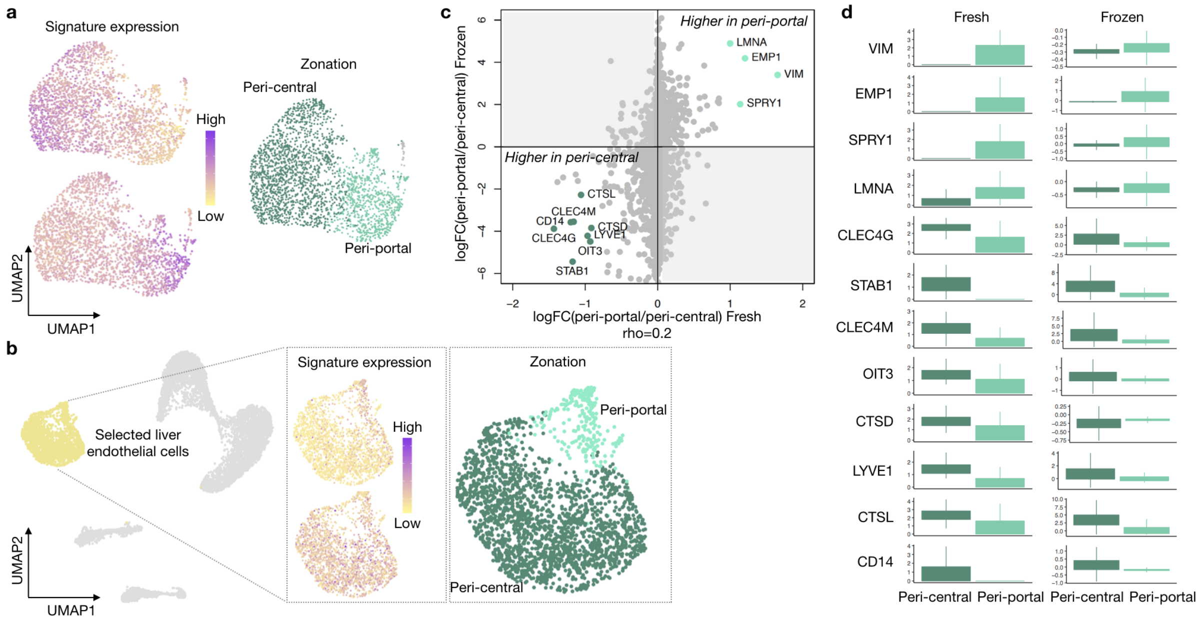


**Extended Data Fig. 4 | Identification of the spatial pattern of zonation in liver sinusoidal endothelial cells (LSEC). a**, UMAP plot of fresh tissue LSEC showing average gene expression of the previously described zonation representing genes for pericentral (upper left) and periportal (lower left) zones and accordingly assigned periportal and pericentral subclusters (right). **b**, UMAP plot of the frozen tissue showing expression signatures of periportal (upper left) and pericentral (lower left) marker genes in LSEC and according to that assigned periportal and pericentral subclusters (right). **c**, Fold change showing differential expression in LSEC from periportal and pericentral subclusters in fresh (x-axis) and frozen (y-axis) tissue data (Spearman's rho=0.2, $p = 4.6 \times 10^{-27}$). Genes sharing the zonation pattern, i.e. genes with abs(logFC)>0.9 for fresh and abs(logFC)>2 for frozen are shown in light green (periportal) and dark green (pericentral), respectively. **d**, Expression of identified genes shown in panel c) across LSEC from periportal and pericentral subclusters in fresh and frozen tissue datasets.

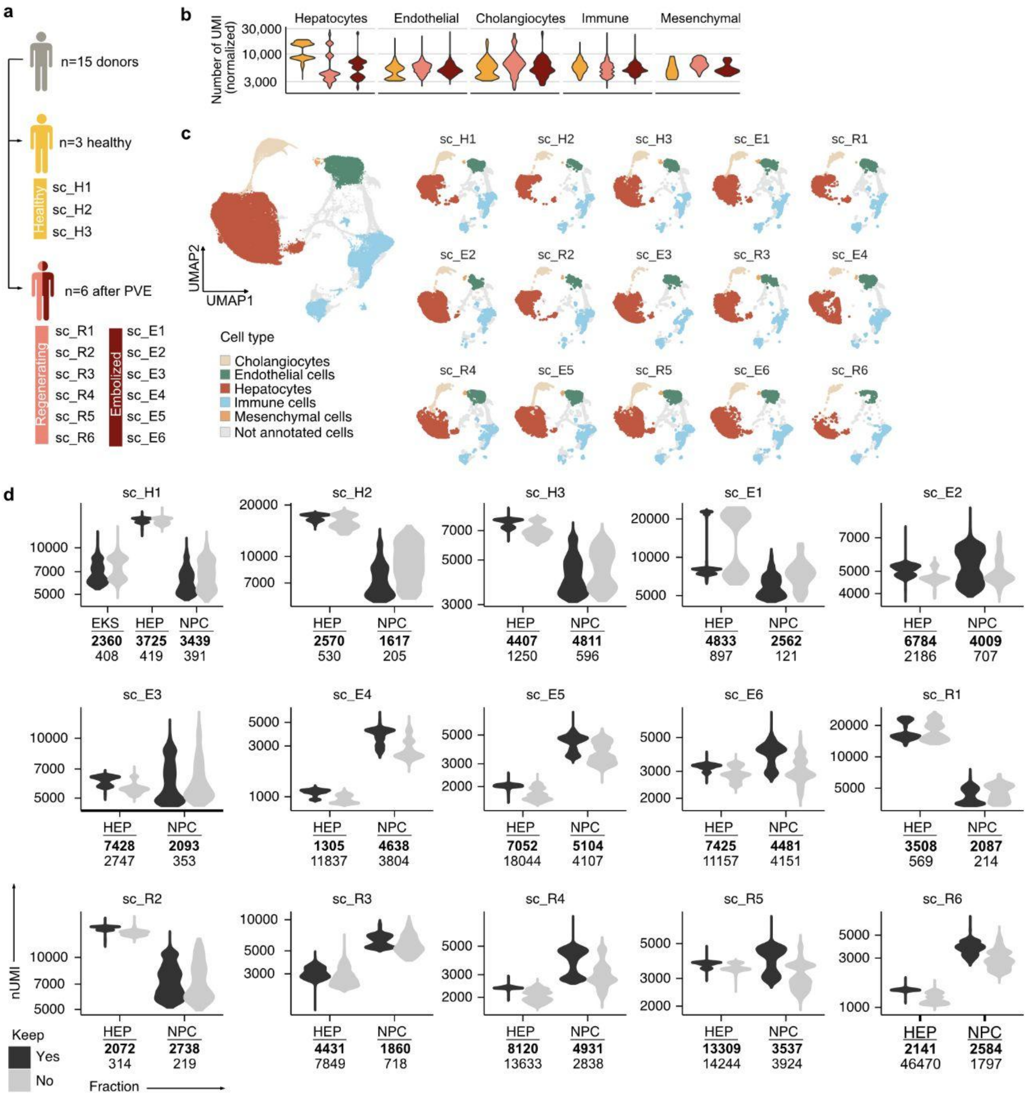


**Extended Data Fig. 5 | Gene expression coverage per donor and condition. a,** Schematic shows information on acquired liver tissues from all donors (top) including healthy (middle, yellow) and after PVE (bottom). From each donor after PVE we obtained regenerating (salmon) and embolized (dark red) conditions. sc_H, single-cell RNA-seq Healthy; sc_R, single-cell RNA-seq Regenerating; sc_E, single-cell RNA-seq Embolized**. b,** Number of detected unique molecular identifiers (UMIs) per annotated cell type**. c**, UMAP plots of scRNA-seq data for patient-derived liver tissues (n=131,961 cells) from all medical conditions and colored by major cell types are shown for all combined medical conditions (left) and for each donor individually (right). **d,** Number of kept (dark gray) and filtered out (light gray) UMIs per analyzed cellular fraction (EKS, mix of Endothelial, Kupffer and Stellate cells; Hep, Hepatocytes; NPC, non-parenchymal cells (see Methods)) across all samples**.**

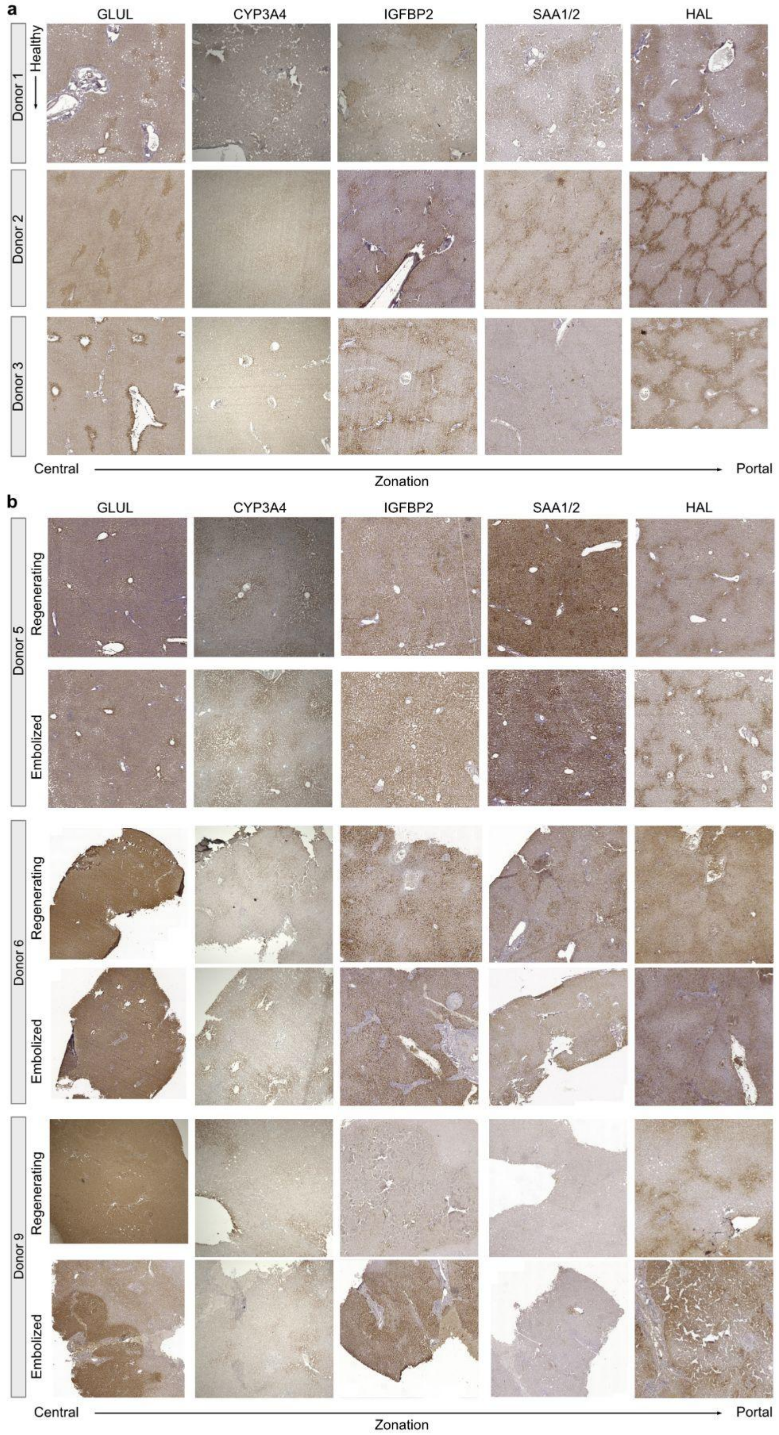


**Extended Data Fig. 6 | Identification of zonation-specific marker expression in healthy and post-PVE liver tissue sections.** Representative sections with DAB stainings of central (GLUL, CYP3A4), middle (IGFBP2) and portal (SAA1/2, HAL) zonation markers in **a**, healthy and **b**, post-PVE (regenerating and embolized) human liver tissues.

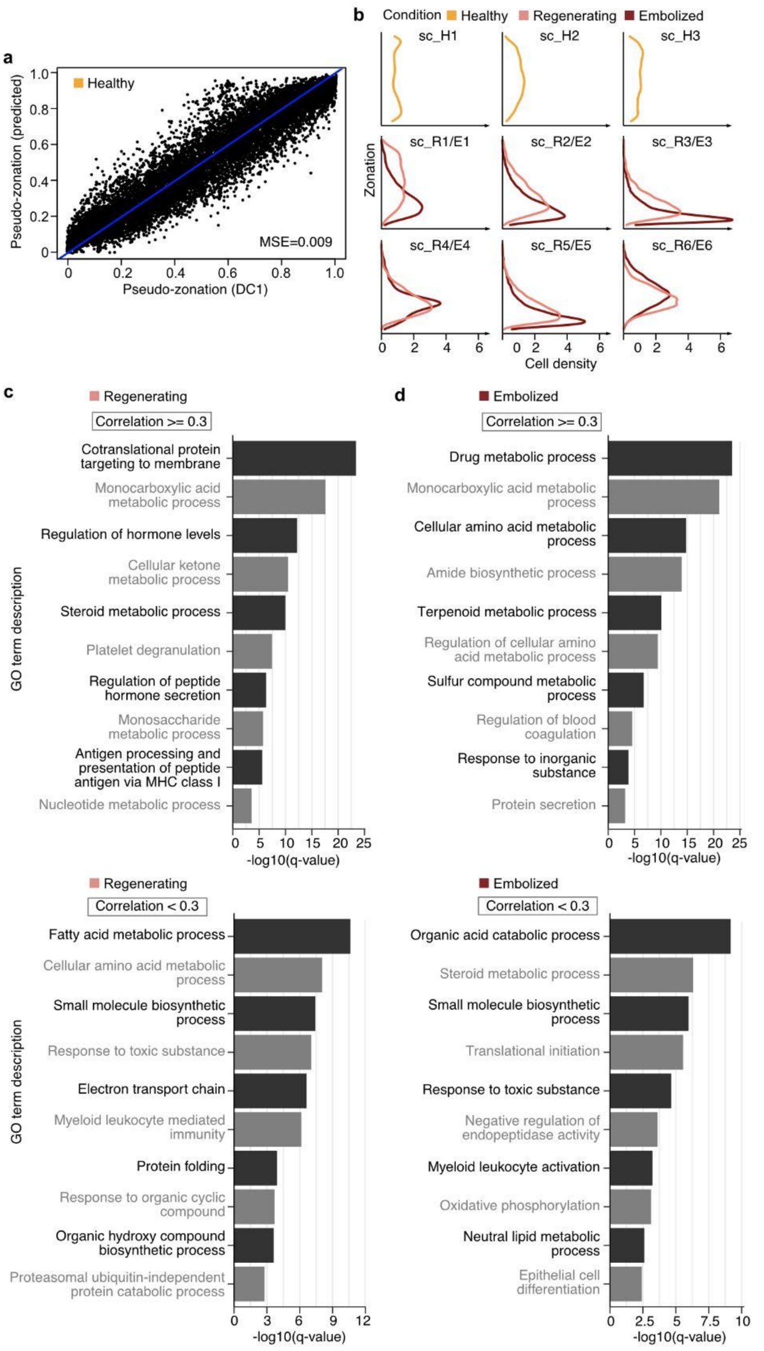


**Extended Data Fig. 7 | Additional information on hepatocyte zonation inference**. **a,** Scatterplot comparing original (x-axis) and predicted (y-axis) hepatocyte pseudozonation. Prediction was done based on 10-fold cross validation. **b,** Cell distribution along zonation in every condition, split by donor. **c, d,** Enriched GO Terms for the genes with fitted expression along pseudozonation correlated (PCC>=0.3, top) and not correlated (PCC<0.3, bottom) between healthy and regenerating (**c**) or embolized (**d**) tissue hepatocytes.

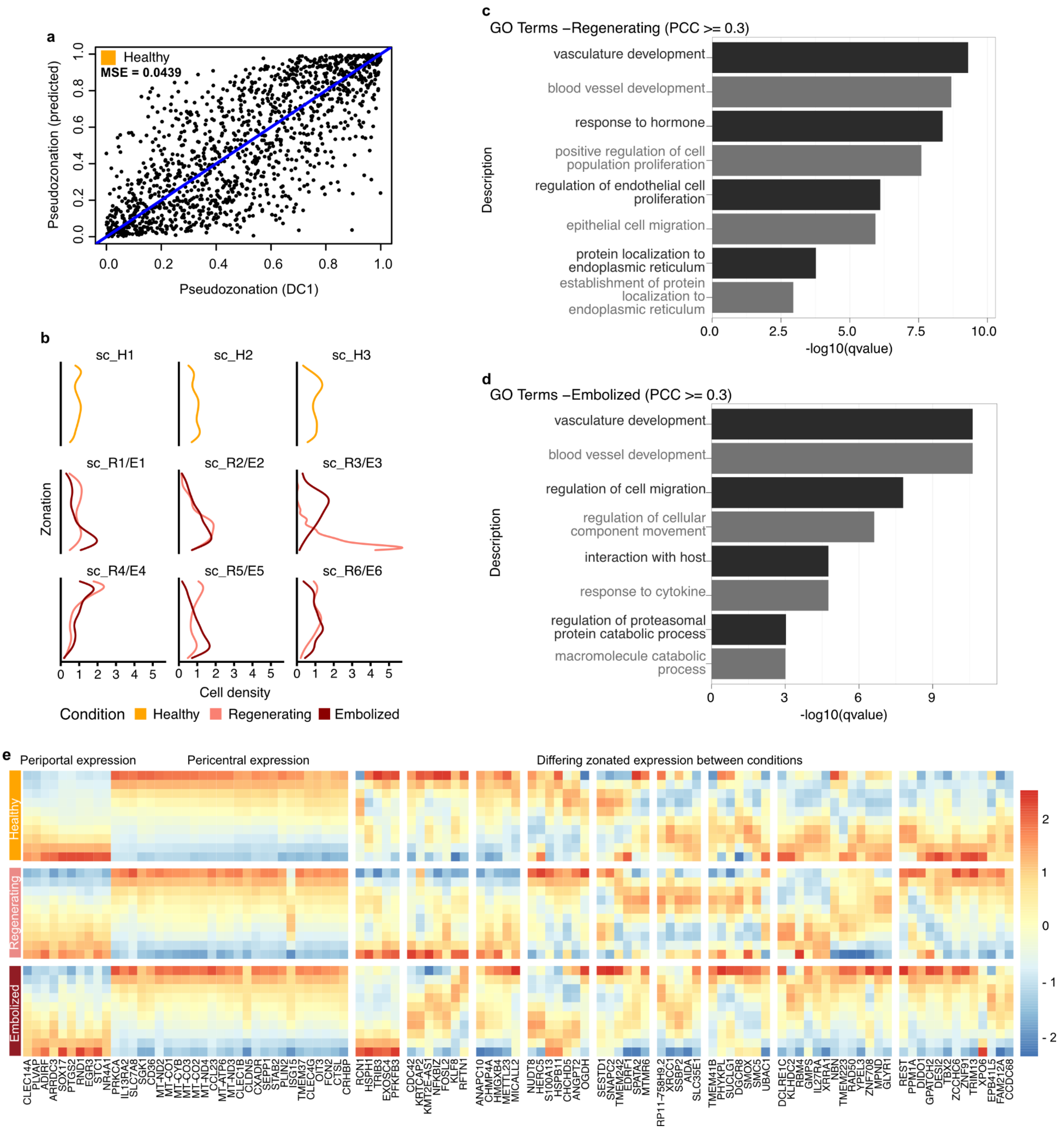


**Extended Data Fig. 8 | Additional information on LSEC zonation inference**. **a,** Scatterplot comparing original (x-axis) and predicted (y-axis) LSEC pseudozonation. Prediction was done based on 10-fold cross validation (see Methods). **b,** LSEC distribution along zonation trajectory in every condition, split by donor. **c, d,** Enriched GO Terms for the genes with fitted expression along pseudozonation correlated (PCC>=0.3, top) between healthy and regenerating (**c**) or embolized (**d**) LSEC. No terms were found enriched from non-correlated genes. **e,** Heatmap shows zonation-specific periportal, midzonal, and pericentral LSEC gene expression, grouped by similarity between conditions. Genes were chosen from the top correlated and anti-correlated genes between healthy and regenerating, and healthy and embolized conditions.

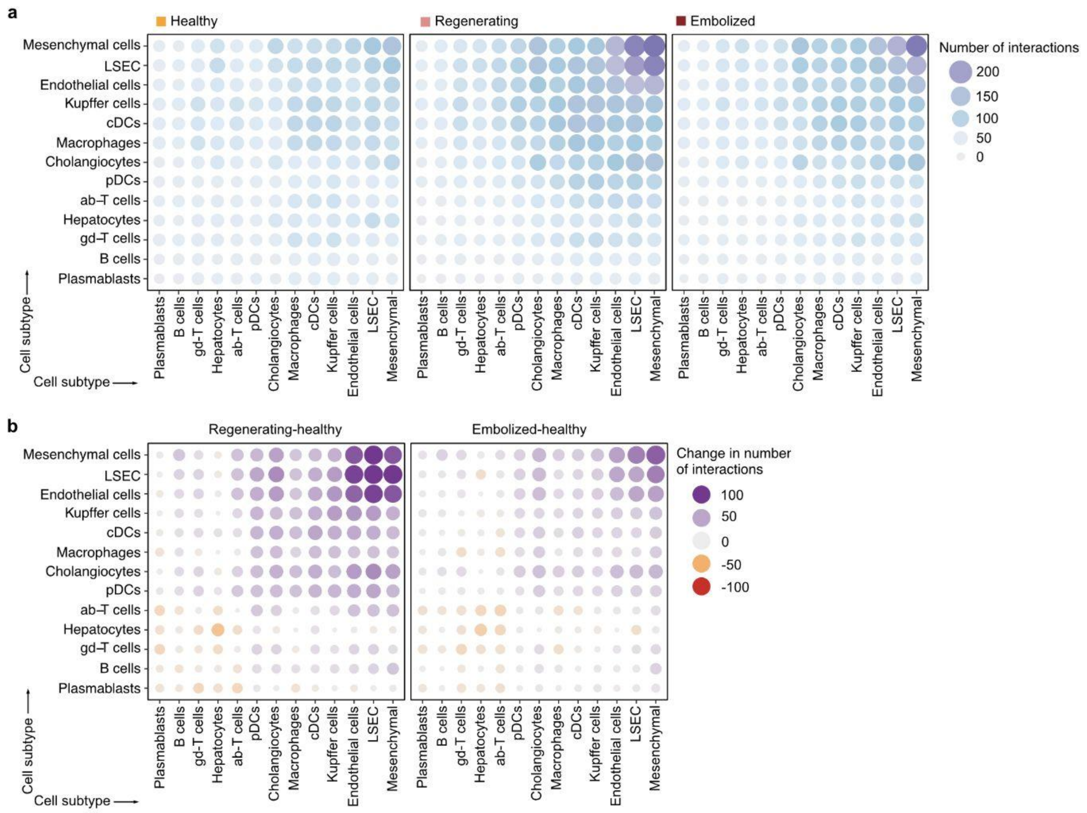


**Extended Data Fig. 9 | Differences in the number of cell-cell interactions between conditions. a**, Absolute number of interactions per condition (healthy, left; regenerating, middle; embolized, right) involving each cell type pair. **b**, Difference in number of interactions for each pair of cell types between healthy and regenerating (left) or embolized (right).

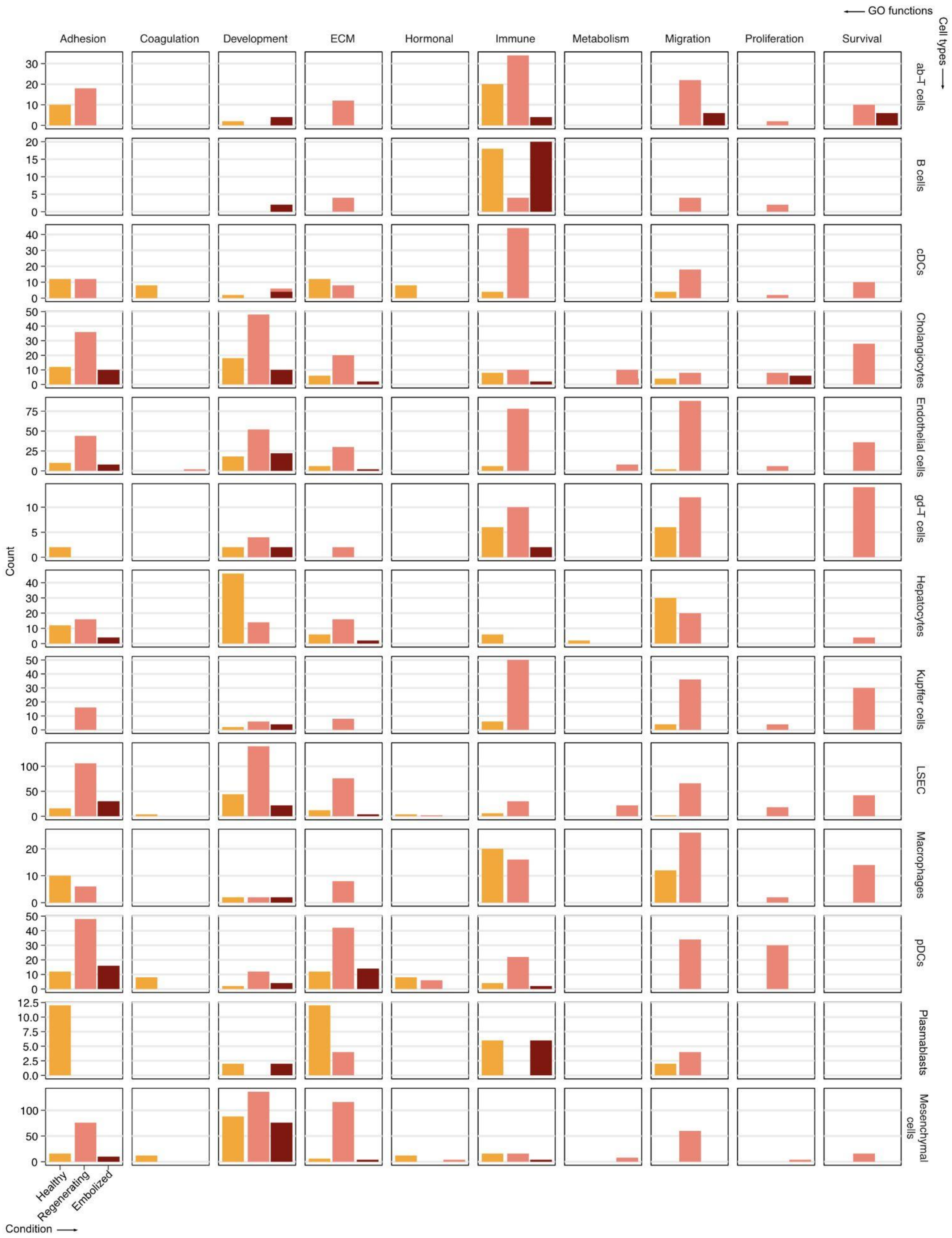


**Extended Data Fig. 10 | Number of unique interactions belonging to each curated function across all pairs of cell types and conditions.** Only interactions differing in at least one condition were considered.
